# Pathogen-driven CRISPR screens identify TREX1 as a regulator of DNA self-sensing during influenza virus infection

**DOI:** 10.1101/2023.02.07.527556

**Authors:** Cason R. King, Yiping Liu, Katherine A. Amato, Grace A. Schaack, Tony Hu, Judith A Smith, Andrew Mehle

## Abstract

Intracellular pathogens interact with host factors, exploiting those that enhance replication while countering those that suppress it. Genetic screens have begun to define the host:pathogen interface and establish a mechanistic basis for host-directed therapies. Yet, limitations of current approaches leave large regions of this interface unexplored. To uncover host factors with pro-pathogen functions, we developed a novel fitness-based screen that queries factors important during the middle-to-late stages of infection. This was achieved by engineering influenza virus to direct the screen by programing dCas9 to modulate host gene expression. A genome-wide screen identified the cytoplasmic DNA exonuclease TREX1 as a potent pro-viral factor. TREX1 normally degrades cytoplasmic DNA to prevent inappropriate innate immune activation by self DNA. Our mechanistic studies revealed that this same process functions during influenza virus infection to enhance replication. Infection triggered release of mitochondrial DNA into the cytoplasm, activating antiviral signaling via cGAS and STING. TREX1 metabolized the mitochondrial DNA preventing its sensing. Collectively, these data show that self-DNA is deployed to amplify host innate sensing during RNA virus infection, a process tempered by TREX1. Moreover, they demonstrate the power and generality of pathogen driven fitness-based screens to pinpoint key host regulators of intracellular pathogens.

## Introduction

Intracellular pathogens depend upon the host cell for replication. They exploit, and in some cases repurpose, cellular components and pathways to promote replication^1^. At the same time, pathogens must evade or suppress innate immune responses deployed by the cell to prevent infection or suppress replication. The balance between these pro- and anti-pathogen forces influences the outcome of an infection, the severity of disease, and even the potential to establish a pandemic outbreak^2,3^.

Influenza virus is a serious public health threat causing annual epidemics and occasional pandemics with significant morbidity and mortality. Identifying cellular genes and proteins required by influenza virus is essential to understanding the viral life cycle and establishing a mechanistic foundation for the development of host-directed anti-viral therapeutics^4^. Diverse systems-level approaches have provided an initial global overview of the interface between influenza virus and its host^5–11^. CRISPR/Cas9-based approaches further accelerated host:pathogen studies^12–14^. Genome-wide CRISPR/Cas9-knockout screens enabled high-throughput functional genomics that identified host dependency factors for influenza A virus^15–18^. CRISPR-activation (CRISPRa), an approach that uses a catalytically inactive Cas9 (dCas9) and engineered sgRNAs to recruit transcriptional activators to targeted genes for gain-of-function screening, has also revealed cellular factors that restrict infection by influenza A and B viruses^19,20^.

Despite the success of these screens, they have been constrained by technical and biological limitations. Most genome-wide screens relied on knockdown or knockout loss-of-function approaches, which only probe those genes already expressed in the system under study and are limited in their ability to detect contributions from genes essential for cell viability, genes with redundant functions, or gene products needed in limited quantities. These approaches almost all manipulated the host prior to infection, skewing the starting population before it was even challenged with virus. Loss-of-function screens also showed a strong bias toward steps involved in entry, repeatedly identifying the same key processes for influenza and other viruses^5–7,13,17,19,20^. Finally, most of the prior approaches measured infection, but not replication *per se*, failing to capture virus:host interactions throughout the entire viral life cycle. Clever variations on classic knockout screens where host-gene target information is encoded within the viral genome have begun to address this by probing more steps of the viral life cycle^21–23^. Due to these limitations, genetic screens have identified only one quarter of the predicted ∼2800 genes in the influenza virus interactome^24^, and even less is known about how those genes impact steps subsequent to entry^25^. New strategies are therefore needed to expand and complement current approaches and reveal a fuller spectrum of host:pathogen interactions^1^.

To address this gap, we developed a new genetic screen where the pathogen programs dCas9 to modulate host gene expression, a technique we call TRPPC (transcriptional regulation by pathogen-programmed Cas9). TRPPC combines viral reverse genetics with CRISPR technology, engineering replication-competent viruses to express sgRNAs that direct CRISPR-activation or -inhibition of targeted host genes. Because sgRNAs are expressed by the virus, screening occurs only in infected cells after viral transcription initiates, focusing on the middle to late stages of infection. We show that TRPPC viruses that activate pro-viral factors gain a replicative advantage and come to quickly dominate the viral population, whereas those that activate anti-viral factors are rapidly lost. This creates a fitness-based screening platform where the pathogen itself does the heavy lifting to pinpoint the most critical cellular regulators of infection. These key advances – where the pathogen drives the screen, relative fitness inherently rank-orders putative co-factors, and gain- and loss-of-function approaches are both possible – sets TRPPC apart from other systems-level approaches. Our genome-wide TRPPC-activation screen identified TREX1 (three prime repair exonuclease 1) as a potent pro-viral factor. Mechanistic studies showed that influenza virus infection triggers release of mitochondrial DNA into the cytoplasm activating the cGAS-STING antiviral pathways. TREX1 degrades this cytoplasmic DNA to temper DNA sensing and innate immune activation, promoting influenza virus replication. Our new screening technology revealed that DNA-sensing pathways activate antiviral responses to the RNA-based influenza virus, and that TREX1 is a cellular factor with inadvertent pro-viral activity that suppresses this innate immune activation.

## Results

### Establishment of transcriptional regulation by influenza-programmed Cas9

To enable discovery of new host factors regulating replication, we developed the pathogen-driven TRPPC screen where the pathogen delivers an sgRNA to program CRISPR-activation (CRISPRa)^26,27^. Pathogen-driven screening requires that the sgRNA be encoded by the pathogen genome. In this way, the pathogen modulates the host to alter its fitness, the sgRNA sequence in the pathogen genome identifies the gene targeted in the host, and the relative abundance of any particular sgRNA in the population after screening is a proxy for the impact of the targeted protein on pathogen replication. We developed TRPPC for influenza A virus (IAV) by exploiting its ability to express non-coding RNAs. The influenza virus *NS* gene encodes NS1 and NS2/NEP in overlapping reading frames. It was previously shown that *NS* can be re-engineered to create a replication-competent viruses with non-overlapping reading frames for NS1 and NS2/NEP that are separated by an artificial intron, a so-called split NS^28^ (Fig 1A). miRNAs encoded in this intron are processed by Drosha without disrupting viral protein expression, genome integrity, or replication^22,28^. We reasoned this system could be used to encode an sgRNA that would be liberated by endogenous miRNA processing enzymes to program CRISPRa, resulting in TRPPCa.

**Fig. 1.**
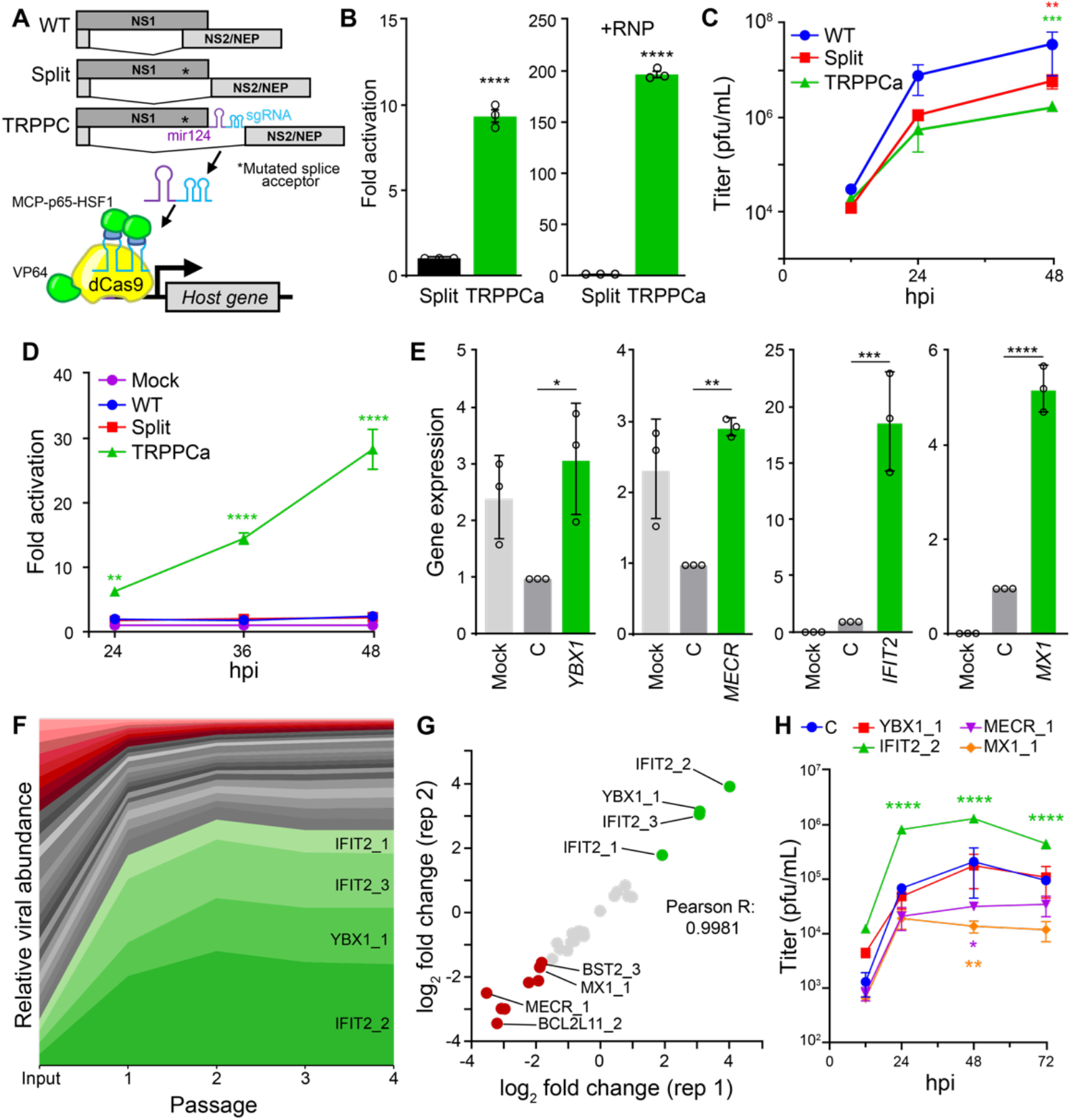
Transcriptional regulation by influenza-programmed Cas9 (TRPPC) manipulates host gene expression to enable fitness-based screening. **A**, Cartoon of re-engineered TRPPC *NS* genome segment and TRPPCa-mediated gene expression. The sgRNA directs VP64-dCas9 to specific genome targets while two MS2 hairpins inserted in the sgRNA recruit MCP-p65-HSF1. **B**, TRPPCa of a luciferase reporter in 293T cells. Cell were transfected with vectors expressing viral genomic RNA for *NS, Split NS* that lacks an sgRNA, or *TRPCC NS* targeting the reporter promoter. Activation was measured in the presence (+RNP, right) or absence (left) of the viral replication machinery. **C**, Multicycle replication of IAV harboring a TRPPC-*NS* segment in A549 cells. **D**, Virally delivered sgRNA activates reporter gene expression in a multicycle infection. A549-CRISPRa cells were inoculated with virus encoding the indicated *NS* segment (MOI = 0.05), or mock treated, and luciferase reporter was measured over the course of infection. **E**, Virally delivered sgRNAs activate expression of host genes from the endogenous locus. A549-CRISPRa cells were inoculated with TRPPC viruses (MOI = 5) targeting the indicated gene, a non-targeting control (C) or mock. Host gene expression was measured at 8 hpi via RT-qPCR. **F**, A pool of 34 TRPPC viruses targeting a collection of 10 potential pro- or antiviral host genes were subject to 4 rounds of selection in A549-CRISPRa cells cells. Viruses present at each stage of selection were quantified by deep-sequencing and normalized sgRNA composition is depicted. Viruses activating proviral genes enriched at least 3-fold are colored green, while viruses activating antiviral genes that are depleted at least 3-fold are colored red. Graph is representative of mean values for 2 replicate screens. **G**, TRPPCa screens are highly reproducible. Comparison of two biological replications shows nearly identical relative enrichment of TRPPC viruses targeting the indicated host genes after 4 rounds of selection. **H**, TRPPC results reflect changes in viral replication. Multicycle replication in A549-CRISPRa cells of individual TRPPC viruses targeting specific host genes (MOI = 0.01). Data are shown as grand mean of 3 replicates ± SEM (B, D) or mean ± s.d. (C, E, F, H). T tests (B), two-way ANOVA with Dunnett’s multiple comparisons test against WT (C, D, H), and one-way ANOVA with Dunnett’s multiple comparisons tests (E) were performed (*p<0.05, **p<0.01, ***p<0.001, ****p<0.0001).

The overlapping reading frames of *NS* were split and the sgRNA was placed within the artificial intron downstream of the primary microRNA-124 (Fig. 1A). miR124 is neuron-specific and its expression by influenza virus in lung cells does not alter viral replication^28^; it is present in our construct solely to direct processing of the intron to release the downstream sgRNA. Production of a functional sgRNA was tested by expressing the *TRPPC NS* gene in cells with the other CRISPRa components. The CRISPRa system we used recruits the transcriptional activator VP64 by fusing it to dCas9. The sgRNA has also been engineered to contains two MS2 hairpins that recruit the transcriptional activators p65 and HSF1 that are fused to the MS2 coat protein^26^. A *TRPPC NS* gene targeting sequence upstream of a minimal promoter in a luciferase reporter construct increased lucifersase expression by almost 10-fold compared to controls (Fig. 1B). We repeated these experiments in cells co-expressing the viral polymerase and nucleoprotein (NP), which replicates and transcribes viral genes, increasing *TRPPC NS* expression and better mimicking an infection. Under these conditions, TRPPCa drove ∼200-fold activation in 293T cells and also activated reporter expression in human lung A549 cells (Fig 1B, S1). A similar design strategy was used to demonstrate TRPPCa for the primary isolate from the 2009 pandemic A/California/07/2009 (CA07) and for influenza B virus (IBV) (Fig S1B). By exchanging activators for the repressive dCas9-KRAB^29^, *TRPPC NS* can also direct inhibition (Fig S1C).

We rescued replication-competent viruses to test TRPPCa during infection. TRPPCa virus generated on the A/Puerto Rico/8/1934 (PR8; H1N1) background replicated to high titers in multiple cell lines, with only modest decreases in overall titers and plaque morphology indistinguishable from WT (Fig. 1C, S1D, S1E). The *TRPPC NS* genome segment was stable through at least four passages of multicycle replication (Fig. S1D). A TRPPCa virus targeting our reporter construct induced luciferase expression, with expression increasing throughout the course of infection (Fig. 1D). TRPPCa viruses targeting endogenous loci also increased gene expression (Fig. 1E). Viruses were generated targeting known pro-(*YBX1, IFIT2)* and antiviral (*MECR, MX1)* genes^15,30–32^. These were used to infect A549 cells stably expressing CRISPRa components (A549-CRIPSRa cells) and relative gene expression was measured. Target genes were specifically up-regulated compared to a non-targeting control virus (Fig. 1E). *YBX1* and *MECR* levels are decreased during normal infection, yet targeting viruses restored expression. The IFN-stimulated genes (ISGs) *IFIT2* and *MX1* are already induced by infection itself, and TRPPCa viruses could push their expression levels even higher. Moreover, MX1 is a potent restriction factor for influenza virus, showing that TRPPCa can force expression of genes that disrupt replication^32^.

The ability of TRPPC to specifically modulate endogenous host genes during infection suggests that TRPPC viruses could affect their own fitness, and thus drive a fitness-based screen. We tested and optimized this approach in a proof-of-principle competition with 34 viruses targeting 11 different genes predicted to promote or inhibit replication. Each gene was targeted with 2-3 unique sgRNAs. Viruses were pooled and used to initiate four sequential rounds of infections in A549-CRISPRa cells. Progeny viruses were collected after each round and the change in abundance of each member was quantified by deep sequencing (Fig 1F). Viruses targeting the known pro-IAV factors *IFIT2* and *YBX1* were rapidly enriched, emerging as clear “winners” after only two rounds of screening. By contrast, viruses targeting antiviral factors like *MX1, MECR*, and *BST2* were depleted from the population. While the PR8 strain used here is mammalian adapted and might be expected to be resistant to MX1, over-expression of MX1 can still restrict replication as reflected in our results^33^. TRPPC Control viruses that trigger apoptosis by inducing BimS and BimL from the *BCL2L11* gene also quickly dropped out. The approach was highly reproducible, with results from two independent screens nearly superimposable (Fig. 1G). Fitness effects during bulk screening were recapitulated for most viruses when tested in isolation (Fig 1H). These effects were dependent on the CRISPRa machinery, as all viruses maintained nearly identical replication in WT cells (Fig. S1G). Thus, TRPPC screening is robust and reproducible, rapidly selecting for and ranking viruses with increased fitness.

### Genome-wide TRPPC screens identify new pro-viral host factors

Because TRPPC relies on expression of sgRNAs by the infecting virus, screening only occurs in infected cells and only after viral gene expression has begun, probing the middle to late stages of replication and avoiding technical bottlenecks of other screens. We performed a genome-wide TRPPC screen to discover host factors regulating IAV replication. A validated CRISPRa sgRNA library was cloned into *TRPPC NS* and used to create a pool of TRPPC viruses, where each human gene was targeted by 3 distinct sgRNAs to ensure efficient activation (Fig. S2A)^26,34^. The resulting population was rich, with over 69,000 unique members, and diverse, with a Shannon’s diversity (H’) of 6.51 (Fig S2B).

Starting from the same library, we performed three independent screens over 5 serial passages (Fig. 2A). Infections were initiated at a low MOI with a large excess of cells ensuring at least 170-fold coverage of the library during the first round, and much higher at later rounds as the richness decreased. Virus was titered and sequenced after each passage. Viral titers increased by almost 2 logs over the course of the experiment while population richness and diversity decreased dramatically, suggesting selection of a more fit population (Fig 2B, S2C). The relative fitness of every virus was gauged by tracking its abundance across each round of competition. In all 3 replicates, more than 100 individual TRPPC viruses were enriched at least 4-fold by the end of the competition (Fig. 2C-E). Enrichment was independent of the abundance of any individual virus in the starting population. In all three screens, very rare members in the starting population became enriched, while very abundant members dropped out (Fig. 2D). In each replicate, the total abundance of winning viruses was approximately 2% in the starting library, but eventually rose to encompass >30% of the population after 5 rounds of screening. Replicate screens were highly reproducible. Spearman’s ρ was between 0.87-0.89 for pairwise comparisons of total populations; for viruses enriched at least 4-fold, 56 gene targets were shared in all 3 replicates and 32 additional genes were shared between two replicates (Fig 2F). sgRNA sequences encoded by the enriched viruses correspond to the cellular genes they activate. To identify these putative pro-viral factors, we performed gene-level analysis of the 3 replicate screens comparing the starting library to the selected viruses after passage 5^35^. This yielded a refined list of enriched gene targets (Fig 2E, 2G). *IFIT3* was amongst our top hits. IFIT3 expression was previously shown to enhance viral replication by promoting viral protein production during the middle to late stages of infection, providing confidence in the ability of TRPPC screening to identify this population of pro-viral factors^15^. Gene enrichment analysis of the top 100 ranked genes did not reveal statistically significant biological process groups. Because our TRPPC screens focused on all infection events post-entry and is otherwise unbiased, a broad list of hits is unsurprising. However, molecular function analysis did uncover modest enrichment for factors involved in ubiquitination pathways (Fig. S2D-E), consistent with the generic pro-viral role of ubiquitination during influenza virus infection^36^. These data establish TRPPC as a powerful, unbiased platform where the pathogen does the work to pinpoint key regulators of replication and inherently rank-order the impact of the implicated host genes.

**Fig. 2.**
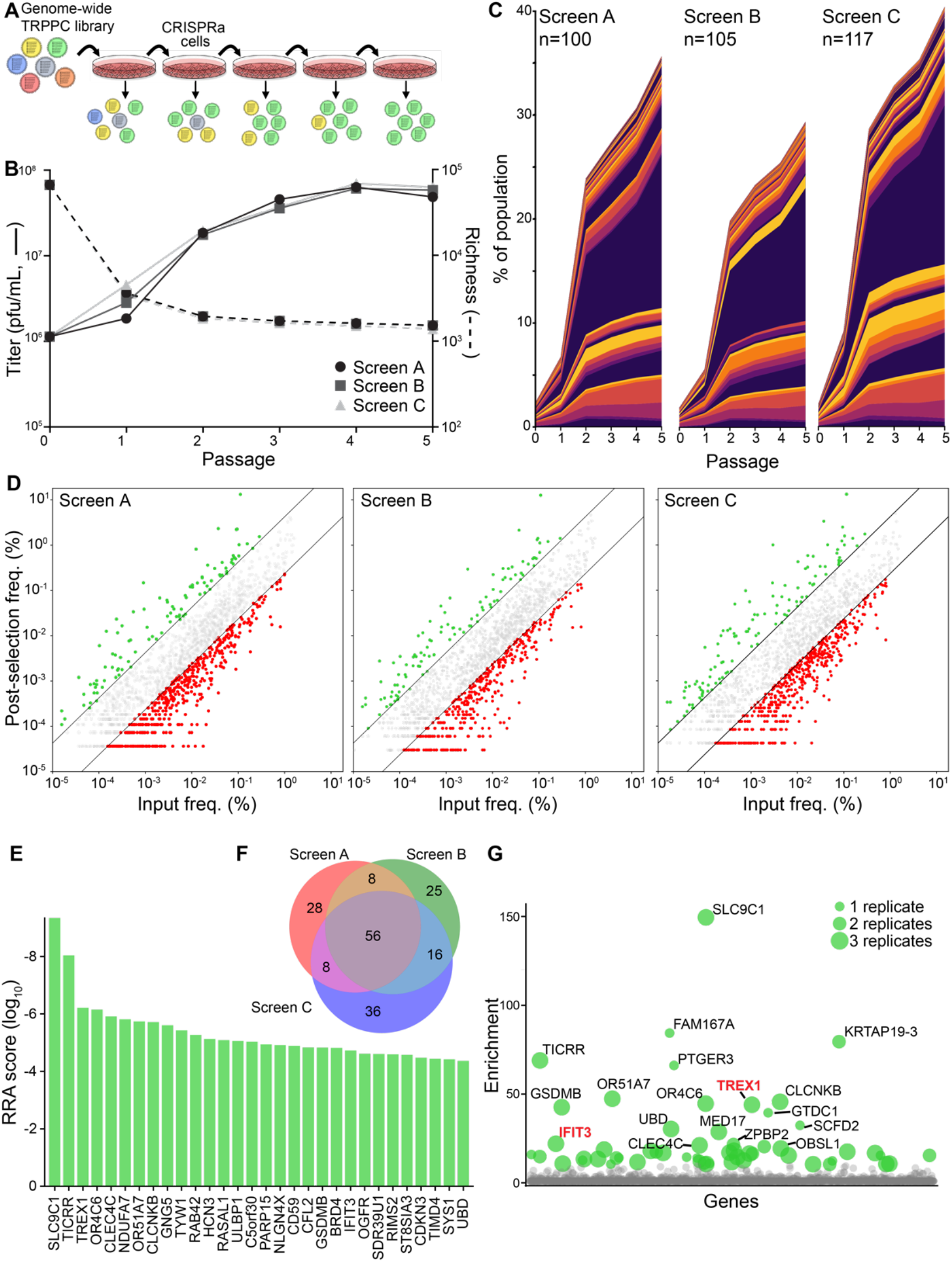
Genome-wide TRPPC screens identify new pro-IAV host factors. **A**, Experimental design of a genome-wide TRPPC screen in CRISPRa cells. **B**, Viral titers (left axis, solid line) and population richness (right axis, dashed line) were measured of 5 sequential rounds of TRPPCa selection. Data for independent screens A, B and C are shown. **C**, Stack plot of the abundance of individual TRPPC viruses in three independent genome-wide screen. Viruses enriched >4-fold at passage 5 are plotted for each replicate, with number of enriched viruses indicated for each screen. Colors are used to distinguish each member, but are not unique to any specific sgRNA. **D**, Final abundance of individual TRPPC viruses at passage 5 as a function of their abundance at passage 0 for all replicates. Colors represent viruses >4-fold enriched (green) or >4-fold depleted (red) or unchanged (grey). **E**, Robust ranking aggregation for top hits. MAGeCK gene scores for the top 30 genes in the TRPPC screens. **F**, Venn diagram of genes enriched >4-fold in the 3 screen replicates. **G**, Bubble plot of positive selection values for all genes in the screen. Bubble size indicates the number of replicate screens in which that gene was detected. Colored dots represent genes >10-fold enriched, with labelled dots representing genes >20-fold enriched.Genes are randomly positioned along the x-axis.

### TREX1 is a pro-viral host factor for RNA viruses

We selected *TREX1* to validate our screen and explore mechanism as it was one of the highest-ranked candidates, was detected in all three screens, and has a defined function^37^. *TREX1* encodes TREX1, a 3’-5’ cytosolic DNA exonuclease that degrades single- and double-stranded DNA^38^. We created three clonal TRPPC viruses each targeting distinct sites in the *TREX1* promoter; these included the same sgRNA enriched in our genome-wide screen along with two new sgRNAs designed with a different rule set^39^. IAV infection itself caused a slight induction of *TREX1*, whereas all *TREX1*-targeting viruses caused a much larger increase in mRNA and protein levels compared to a non-targeting virus (Fig 3A). The increased induction of TREX1 was dependent on the CRISPRa machinery, as it was absent in WT A549 cells (Fig S3A). *TREX1*-targeting TRPPC viruses also replicated faster and to higher levels than a non-targeting control in A549-CRISPRa cells (Fig. 3B), but not WT cells (Fig S3B). *TREX1*-targeting and control viruses were pooled and used to initiate a competitive infection (Fig 3C). *TREX1*-targeting viruses were significantly enriched by the end of the infection, almost eliminating non-targeting viruses and a virus activating the antiviral *MX1* gene. Together these data validate TREX1 as a top hit from our screen and further demonstrate the ability of TRPPC to act as both a discovery and validation platform of important virus-host interactions.

**Fig. 3.**
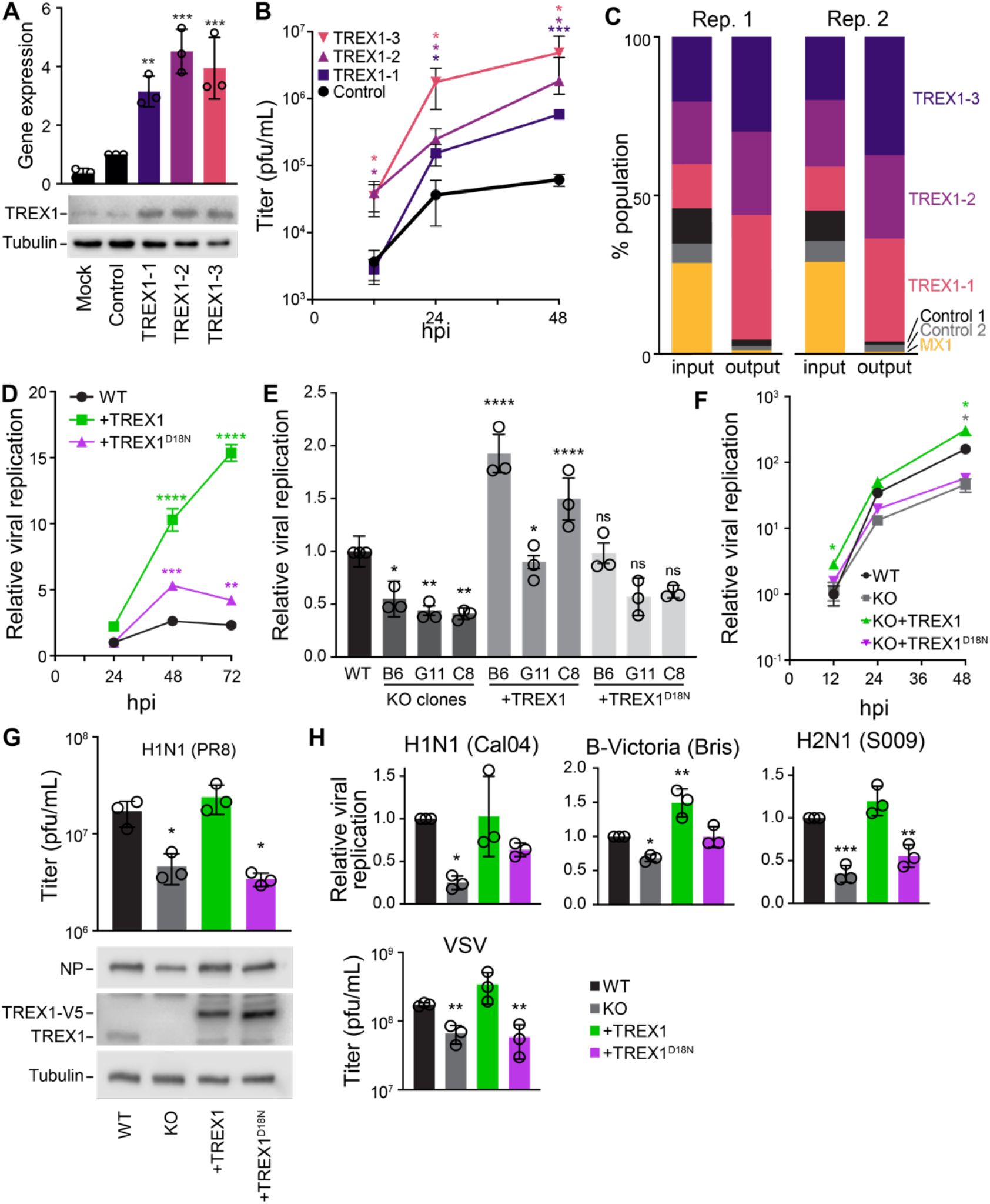
The 3′-5′ DNA exonuclease TREX1 is a pro-viral host factor for RNA viruses. **A**, Multiple TRPPC viruses with distinct targeting sequences activate *TREX1* expression. A549-CRISPRa cells were inoculated (MOI = 1) with viruses targeting different sites in the *TREX1* promoter or a non-targeting control. TREX1 expression was measured relative to mock by RT-qPCR at 10 hpi and western blotting at 12 hpi. **B**, Multicycle replication of *TREX1*- or non-targeting TRPPC viruses in A549-CRISPRa cells (MOI = 0.01). Titers determined by plaque assay. **C**, A pool of TRPPC viruses were competed for 48 h during replication in A549-CRISPRa cells (pooled MOI = 0.05). Relative abundances at the start (input) and end (output) of the infection for each virus was determined by sequencing and shown for 2 independent replicates. **D**, Multicycle replication of a WSN influenza A reporter virus in WT A549 cells or lines stably expressing TREX1 or the catalytic mutant TREX1^D18N^. Replication was normalized to viral titers in WT cells at 24 hpi. **E**, Viral replication was measure at 48 hpi (MOI = 0.05) in 3 distinct TREX1-KO clones inoculated with a WSN influenza A reporter virus. Clones were complemented with TREX1 or TREX1^D18N^, where indicated. Values are relative to replication in parental WT A549 cells. For statistical analyses, KO clones were compared to WT, whereas complemented clones were compared to the matched KO. **F**, Multicycle replication of a WSN influenza A reporter virus (MOI = 0.05) in WT A549 cells, TREX1-KO cells, or complemented cell lines. Values are compared to replication in parental WT A549 cells. **G**. Viral titers at 48 hpi (above) and NP protein levels at 24 hpi (below) in cells inoculated with PR8 (MOI = 0.01) **H**, Replication of reporter influenza viruses based on the primary viral isolates CA04 (MOI = 0.5), S009 (MOI = 0.05), B/Bris (MOI = 0.2) at 48 hpi or VSV (MOI = 0.001) at 24 hpi. Values are compared to replication in parental WT A549 cells. Data are shown as grand mean for 3 replicates ± SEM (D-F, H) or mean ± s.d. (A-B, D, G). One-way ANOVA with post-hoc Dunnett’s tests were performed except for (E), which used a one-way ANOVA with post-hoc Tukey’s test (*p<0.05, **p<0.01, ***p<0.001, ****p<0.0001, ns = not significant).

As an orthogonal validation approach, we expressed TREX1 in cells prior to infection. IAV replicated to higher levels in cells stably expressing TREX1 compared to WT controls (Fig 3D). In parallel, we used a TREX1 catalytic mutant (TREX1^D18N^) that results in a loss of function and is a genetic cause of the autoimmune diseases Aicardi-Goutières syndrome (AGS) and familial chilblain lupus^40–42^. The pro-viral activity of TREX1 was dependent on its nuclease activity, as cells expressing TREX1^D18N^ only marginally increases IAV replication (Fig 3D). Similar results were observed when TREX1 was transiently expressed (Fig S3C).

We also used a loss-of-function approach to assess the impact of TREX1. Pooled TREX1-knockout cells supported lower levels of replication for the IAV strain A/WSN/33 (WSN; Fig S3D). Three different clonal *TREX1*^*-/-*^ lines also revealed decreased replication compared to parental WT cells (Fig 3E, S3E-F). We complemented these cells by stably expressing TREX1 or TREX1^D18N^. TREX1 expression restored IAV replication, whereas TREX1^D18N^ did not, confirming that the knockout phenotype was caused by the loss of TREX1 and solidifying the importance of TREX1 catalytic activity (Fig 3E, S3F). Clone C8 was used for all downstream experiments, hereafter referred to as TREX1-KO. Multicycle replication experiments showed that IAV replicates to higher titers at all time points in WT cells versus TREX1-KO cells (Fig 3F). As before, this defect was rescued by stably expressing TREX1, but not TREX1^D18N^ (Fig 3F, S3G). We tested the universality of our observations by measuring replication of various isolates of influenza A or B virus. Our TRPPC viruses were constructed in the PR8 strain. As such, normal levels of PR8 replication and NP expression were dependent on the presence of WT, but not mutant, TREX1 (Fig 3G). Similar results were detected when measuring replication of the 2009 pandemic isolate A/California/04/2009 (CA04; H1N1), an IAV with avianized RNP components from A/Green-winged Teal/Ohio/175/1986 (S009), or the primary influenza B isolate B/Brisbane/60/2008 (Fig 3H). Lastly, this trend also extended to an unrelated rhabdovirus, vesicular stomatitis virus (VSV). VSV replication was suppressed in TREX1-KO cells, but was rescued by expression of WT TREX1 (Fig 3H). Combined, these data demonstrate that the catalytic activity of TREX1 enhances replication of divergent negative-strand RNA viruses.

### DNA-sensing pathways regulate RNA virus replication

The nuclease activity of TREX1 antagonizes cytosolic DNA sensing and innate immune activation, and its dysfunction can cause autoimmune disorders^43–46^. We therefore sought to test whether TREX1 suppresses innate immune sensing during IAV infection, and if this is related to its pro-viral phenotype. We established IFN-stimulated response element (ISRE) reporter cells line in both WT and TREX1-KO A549 cells. WT and TREX1-KO cells were equally responsive to ectopic IFNβ treatment, indicating an intact IFN signaling cascade (Fig S4A). They displayed no difference in ISRE induction when transfected with poly(I:C), a potent dsRNA analog that activates cytoplasmic RNA sensors (Fig S4B)^47^. Further, basal ISRE induction was indistinguishable between WT and TREX1-KO cells (Fig 4A). However, ISRE induction by foreign DNA was dramatically increased in TREX1-KO cells. Transfection of salmon sperm DNA caused a dose-dependent ISRE induction, which was significantly greater in TREX1-KO cells compared to WT (Fig 4A). Loss of TREX1 also dramatically upregulated expression of the endogenous ISGs *IFNB1, IFIT2*, and *MX1* in response to salmon sperm DNA transfection (Fig 4B). Complementing the knockout cells with TREX1 reduced activation of the endogenous ISGs, in some cases below the levels in WT cells (Fig 4B). Thus, loss of TREX1 results in a more potent IFN response to DNA stimulation, including antiviral genes.

**Fig. 4.**
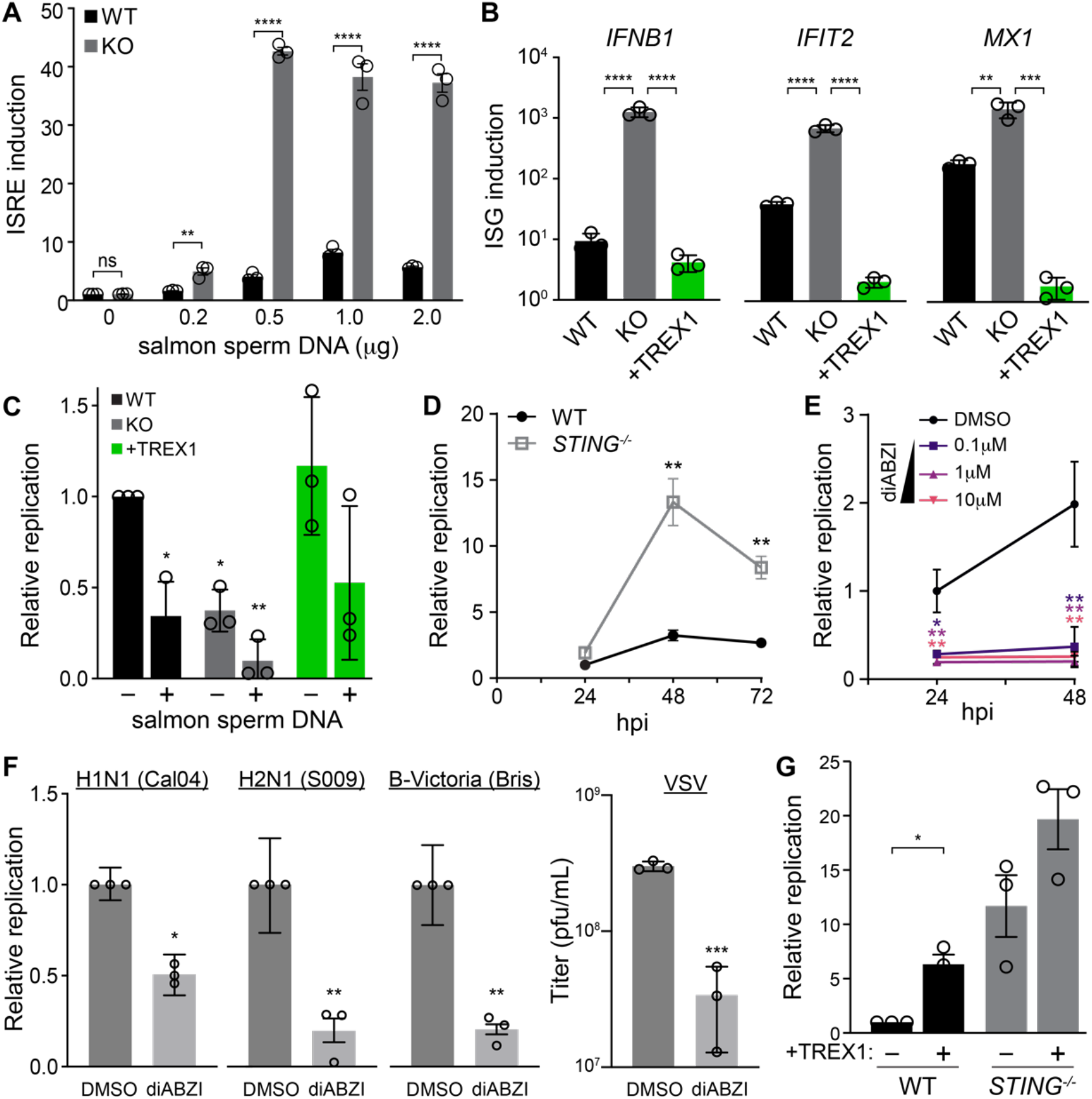
TREX1 regulates RNA virus replication by moderating DNA sensing. **A**, WT and TREX1-KO cells were transfected with the indicated amounts of salmon sperm DNA and innate immune activation was measure with an IFN-stimulated response element (ISRE) reporter. Values are normalized to untransfected WT cells. **B**, WT, TREX-KO, or complemented cells were transfected with salmon sperm DNA. ISG expression relative to mock-transfected cells was measured by RT-qPCR. **C**, WT, TREX-KO, or complemented cells were transfected with salmon sperm DNA prior to inoculation with IAV (MOI = 0.05). Replication was measured at 24 hpi and normalized to mock-transfected WT cells. **D**, Multicycle replication of IAV (MOI = 0.05) in WT or STING-KO A549 cells. Replication is normalized to WT cells at 24 hpi. **E**, Multicycle replication of IAV WSN reporter virus (MOI = 0.05) in A549 cells treated with a STING agonist (diABZI) or a DMSO control. Replication is normalized to DMSO-treated cells at 24 hpi. **F**, Replication of reporter influenza viruses based on the primary viral isolates CA04 (MOI = 0.5), S009 (MOI = 0.05), B/Bris (MOI = 0.2), or VSV (MOI = 0.001) in A549 cells treated with 1μM diABZI or control. Relative replication was measured at 48 hpi for influenza viruses and 24 hpi for VSV. **G**, Replication of an influenza reporter virus (MOI= 0.05) at 48 hpi in WT and STING-KO A549 cells stably expressing TREX1 where indicated. Data are shown as grand mean of 3 replicates ± SEM (A, C-G) or mean ± s.d. (B). Pairwise T tests (A, D, F, G) or one-way ANOVA with post-hoc Dunnett’s tests (B, C, E) were performed (*p<0.05, **p<0.01, ***p<0.001, ****p<0.0001, ns = not significant). Comparisons in C are to untreated WT cells, while those in E are to the DMSO control.

Activating innate immune responses with salmon sperm DNA severely inhibited IAV replication (Fig S4C). This DNA-mediated impairment of replication was further exacerbated in TREX1-KO cells, but replication was partially restored in cells complemented with TREX1 (Fig. 4C). TREX1 metabolizes ligands of cytoplasmic DNA sensors and suppresses their activation, especially the cGAS-STING (cyclic GMP-AMP synthase-stimulator of interferon genes) pathway^48^. We therefore investigated whether this pathway regulated IAV infection. We blocked activation by knocking out *STING*. IAV replicated to much higher levels in *STING*^*-/-*^ A549 cells compared to matched controls (Fig 4D). This increase was comparable to the enhanced replication observed in *MAVS*^*-/-*^ cells that are defective in sensing foreign RNA, the major innate response to IAV^49^ (Fig S4D). Conversely, we artificially activated STING with the chemical agonist diABZI. Prophylactic treatment of cells with diABZI inhibited IAV replication at all concentrations tested (Fig 4E). diABZI treatment also inhibited replication of a larger panel of RNA viruses including CA04, S009, IBV, and VSV (Fig. 4F), as well as SARS-CoV-2^50^. We then tested genetic interactions between TREX1 and STING. Over-expressing TREX1 in WT cells resulted in a ∼6-fold increase in replication, paralleling earlier results (Fig 4G). Over-expressing TREX1 in *STING*^*-/-*^ cells also increased replication, but by less than 2-fold compared to parental *STING*^*-/-*^ cells. Thus, TREX1 regulates flux through the cGAS-STING pathway, and possibly other cytoplasmic DNA sensors^51^. Together, these data reveal that infection by RNA viruses triggers DNA sensing that activates innate immune responses to suppress replication.

### TREX1 degrades self-DNA released during IAV infection

IAV is an RNA virus, and its genome does not have a DNA intermediate stage, raising questions as to how the cGAS-STING pathway is activated. We infected WT, TREX1-KO, and complemented A549 cells with IAV and stained for the presence of dsDNA (Fig. 5A). Puncta of dsDNA were detected in the cytoplasm of cells at 8 hpi in all three cell lines when they were infected, but not in mock treated conditions. TREX1 knockout results in precocious detection of dsDNA as early as 4 hpi, with a higher abundance of puncta throughout. Complementing cells with TREX1 restored a pattern that was similar to WT cells. Puncta were detected in NP-negative cells, suggesting the signal causing cytoplasmic DNA release may not be cell autonomous, or that incomplete infections that lack NP are still capable of initiating this process^52^. A potential source of immunogenic DNA during IAV infection is from host mitochondria^53–55^. We used qPCR to identify and quantify mitochondrial DNA (mtDNA) in cytoplasmic extracts. Infection releases mtDNA into the cytoplasm of human lung cells (Fig 5B), consistent with similar experiments in 293T cells^53^. mtDNA accumulated to higher levels in TREX1-KO cells, but was restored to WT levels in complemented cells (Fig 5B), reinforcing our immunofluorescence results. These data confirm that IAV triggers the release of mtDNA into the cytoplasm and now show that TREX1 regulates accumulation of this potentially immunogenic DNA.

**Fig. 5.**
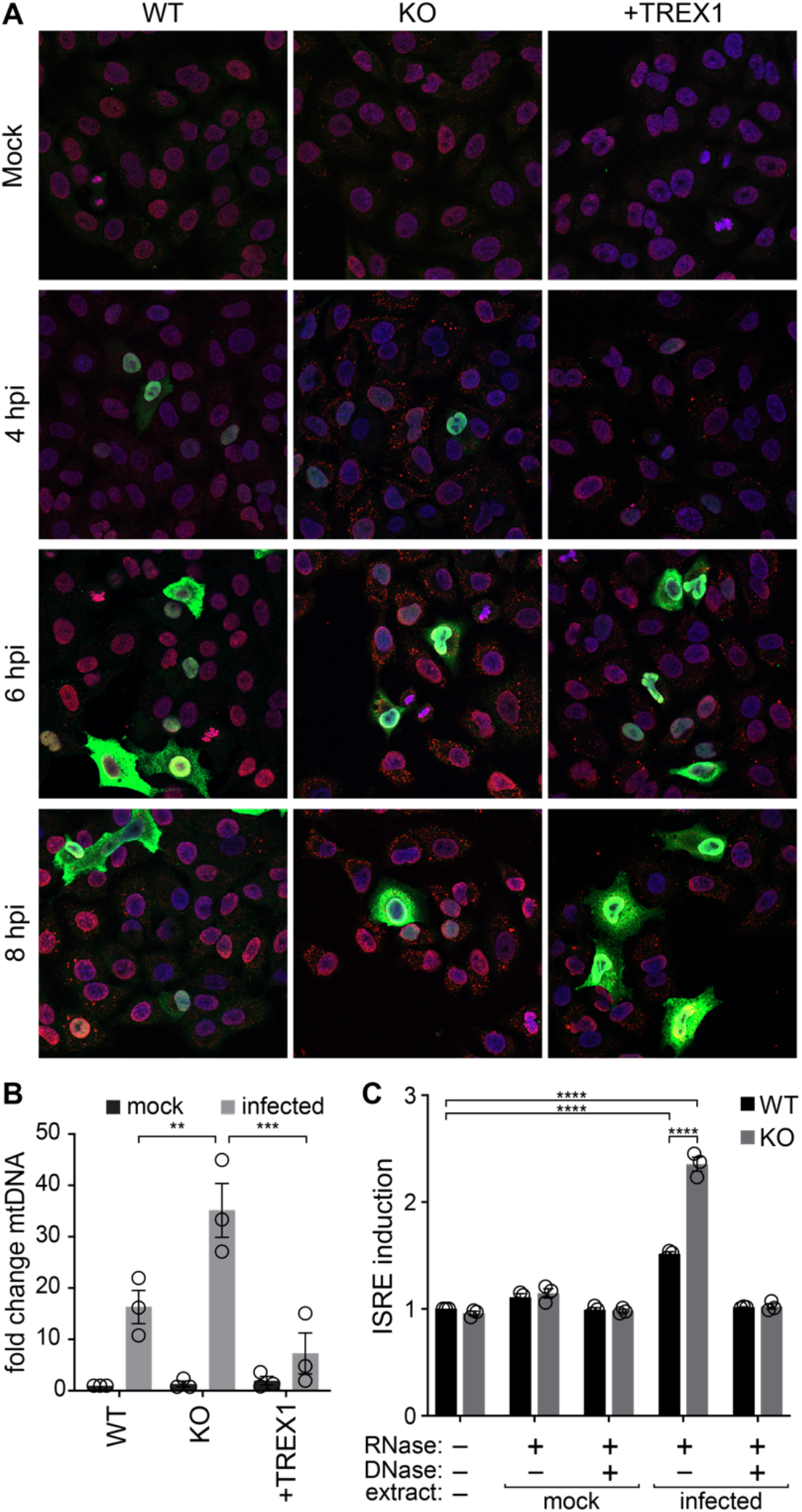
TREX1 degrades self-DNA released during IAV infection. **A**, IAV infection releases dsDNA into the cytoplasm. Immunofluorescence staining of WT, TREX1-KO and complemented A549 cells inoculated with IAV (MOI = 1). Blue = DAPI (nucleus), green = viral NP, red = dsDNA. **B**, Cytosolic extracts were prepared from WT, TREX-KO and complemented A549 cells inoculated with influenza virus (MOI = 1) or mock treated. mtDNA in the cytoplasm was quantified by qPCR and shown relative to mock-infected WT cells. **C**, Cytosolic extracts were prepared from mock or infected A549 cells and re-introduced into WT or TREX1-KO ISRE reporter cells. Where indicated, extracts were pre-treated with nucleases prior to transfection. ISRE activation is normalized to untransfected WT cells. Data are shown as grand mean of 3 replicates ± SEM (C) or mean ± s.d. (B). Two-way ANOVA with Šdák’s multiple comparisons test (B) or a two-way ANOVA with post-hoc Tukey’s tests were performed (**p<0.01, ***p<0.001, ****p<0.0001, ns = not significant).

To test if this host-derived DNA released into the cytoplasm during IAV infection is immunogenic, we permeabilized infected cells to obtain cytoplasmic extracts, recovered nucleic acids from the extracts, and transfected them into ISRE reporter cell lines generated in a WT or *TREX1*^*-/-*^ background. Total nucleic acids from infected cells, which include highly immunogenic viral RNAs^56,57^, caused strong innate immune activation (Fig S5). Extracts were therefore treated with RNase to focus solely on immunogenic DNA. DNA extracted from the cytoplasm of infected cells, but not mock-infected cells, activated the ISRE (Fig 5C). ISRE activation was further exacerbated in TREX1-KO cells. Activation was reversed in all cell types when extracts were additionally treated with DNase, indicating that DNA is the sole immunogenic ligand. Thus, cells lacking TREX1 contain more mtDNA in their cytoplasm and are more sensitive to this DNA. While viral RNA remains the dominant nucleic acid activator, these data show host DNA is also a *bona fide* ligand amplifying antiviral responses via a parallel signaling cascade.

### TREX1 modulates the anti-viral host response to IAV infection

TREX1 enhances IAV replication (Fig 3) and controls the abundance and sensing of mtDNA in the cytoplasm (Fig 5). This raises the possibility that sensing of self-DNA may amplify innate immune responses to IAV. We established a more sensitive nucleic acid sensing assay by extracting total nucleic acids from WT human lung cells infected with IAV. Transfection of untreated nucleic acids into our ISRE reporter cells lines produced an almost 100-fold activation, independent of the presence of TREX1 (Fig 6A). RNase treatment revealed the presence of immunogenic DNA ligands. Again, DNA-dependent activation was significantly stronger when extracts were introduced into TREX1-KO reporter cells compared to WT (Fig 6A, 4A and 5C). Treatment with both RNase and DNase returned ISRE activity to baseline levels, confirming that cells are sensing nucleic acids. Loss of TREX1 also sensitized activation of endogenous ISGs (Fig 6B). *IFIT2* and *MX1* demonstrated DNA-dependent activation in TREX1-KO cells that was suppressed when these cells were complemented with TREX1. Multiple lines of evidence show that IAV infection releases immunogenic self-DNA and TREX1 negatively regulates its detection.

**Fig. 6.**
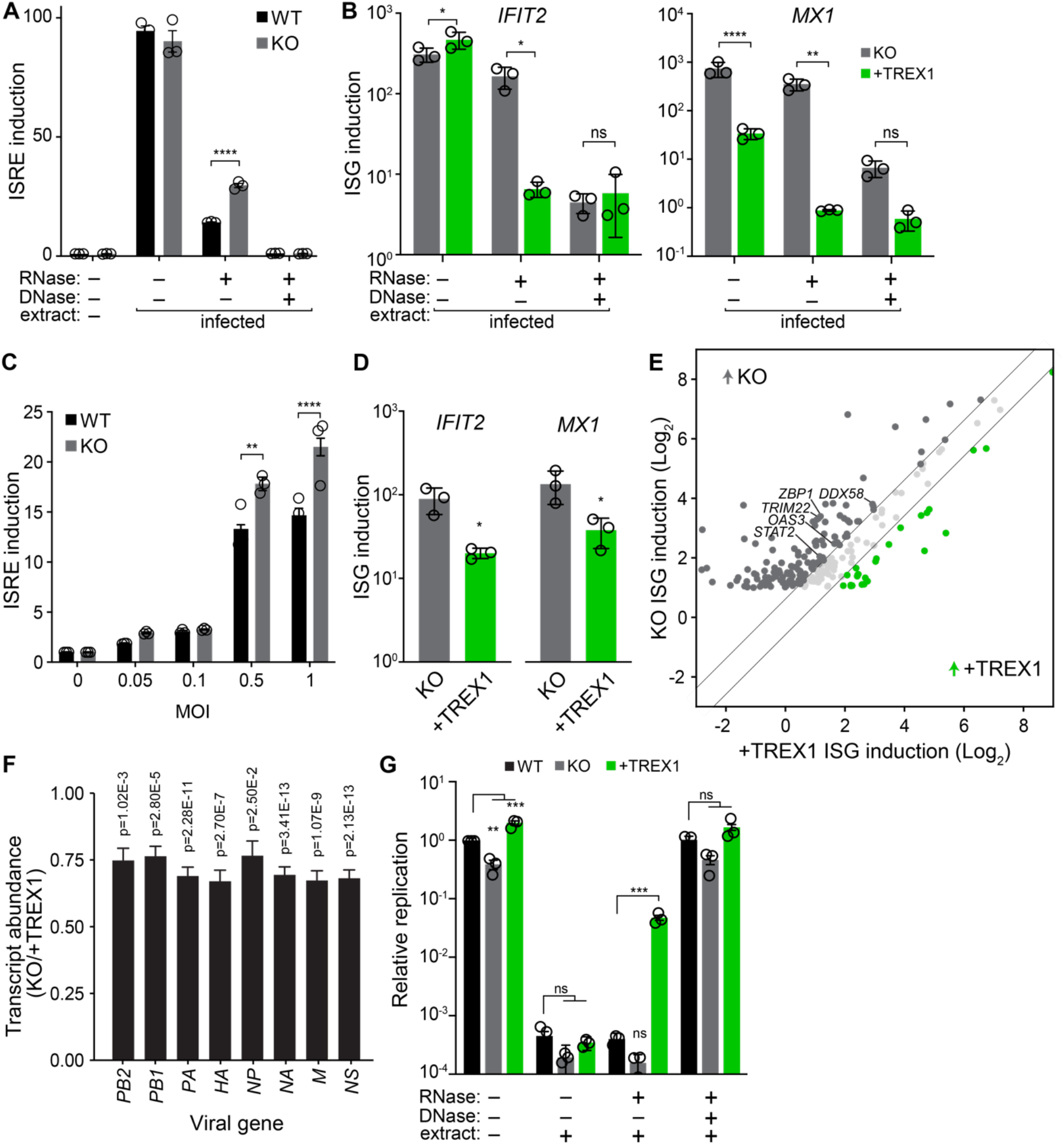
TREX1 tempers the anti-viral host response to influenza virus infection. **A**, ISRE induction was measured in WT and TREX1-KO reporter cells transfected with nucleic acids extracted from infected A549 cells. Extracts were treated with the indicated nucleases prior to transfection. ISRE induction was normalized to untransfected WT cells. **B**, Activation of endogenous ISGs in TREX-KO or complemented cell lines transfected with nucleic acids extracted from infected A549 cells was measured by RT-qPCR. Extracts were treated with the indicated nucleases prior to transfection. **C**, ISRE induction was measured in WT and TREX1-KO reporter cells infected with IAV at the indicated MOIs. Data are normalized to uninfected cells for each cell type. **D**, Activation of endogenous ISGs in TREX1-KO and complemented cells infected with IAV was measured by qRT-PCR. **E**, Comparison of ISG induction (infected/mock) in TREX1-KO and complemented cells. Only ISGs induced ≥2-fold during infection in TREX1 KO cells are shown. Diagonal lines separate ISGs whose induction levels change by at least 50% different between cell lines. **F**, Differential gene expression of IAV transcripts in TREX1-KO versus complemented cells. **G**, IAV reporter replication in WT, TREX1-KO, and complemented A549 cells pre-treated with nucleic acids extracted from infected A549 cells. Extracts were treated with the indicated nucleases. Viral titers were measured 48 hpi and are shown relative to untreated WT cells. Data are shown as grand mean of 3 replicates ± SEM (A, C, G), mean ± s.d. (B, D), or fold change of three independent RNA-seq experiments (E-F). Significance was tested with a two-way ANOVA with Šídák’s mulitple comparisons (A-C), an unpaired T-test (D), a one-way ANOVA within each group with Dunnet’s correction (G), or Wald statistic (F). FDR-adjusted p-values are shown in F. For all others, *p<0.05, **p<0.01, ****p<0.0001, ns = not significant.

Given that TREX1 tempers innate immune activation, we sought to determine if this was the mechanism by which TREX1 enhanced viral replication. WT and TREX1-KO cells showed nearly identical ISRE activity at baseline in untreated conditions (Fig 4A, 5C, 6A). Yet, TREX1-KO cells displayed higher ISRE induction during infection compared to WT (Fig 6C). Expression levels of endogenous ISGs in response to infection were also controlled by TREX1; TREX1-complemented cells showed reduced *IFIT2* and *MX1* expression relative to TREX1-KO cells (Fig 6D). To obtain a comprehensive view of the impact of TREX1 on the innate immune response during IAV infection, we performed RNA sequencing of TREX1-KO or complemented cells that were either mock treated or infected with IAV. Differential gene expression analysis reveals upregulation of multiple GO biological processes involved in antiviral responses in infected cells, regardless of TREX1 expression (Fig S6B). However, when we consider the magnitude of ISG induction, TREX1-KO cells exhibit a much more potent antiviral response. Of the 222 ISGs that exhibited at least a 2-fold change during infection, 133 were expressed higher in TREX1-KO cells compared to TREX1-complemented cells, while only 26 were higher in complemented cells (Fig 6E). These transcript-level data corroborate our results with ISRE reporter cells (Fig 6C). ISGs induced at higher levels in TREX1-KO cells showed strong enrichment for antiviral processes (Fig S6C) and included *DDX58*(RIG-I), *TRIM22, OAS3, STAT2*, and *ZBP-1/DAI*, genes known to be expressed upon STING activation and to suppress IAV replication^50,58–62^. Analysis of viral mRNAs revealed a ∼25% reduction in TREX1-KO cells compared to the complemented cells (Fig 6F), reflecting the higher antiviral state in these cells during infection. This correlates with reduced viral replication in TREX1-KO cells compared to WT or complemented cells (Fig 3E-H). Host gene expression in uninfected TREX1-KO1 cells show very modest enrichment of regulatory pathways of inflammation (Fig 6D), but no dramatic upregulation of antiviral genes.

TREX1 moderates the antiviral IFN response, predicting that sensing of self-DNA should suppress IAV replication. We tested this by transfecting cells with nucleic acid extracts prior to infection. Mock-transfected cells mirrored earlier results, where IAV replicated poorly in TREX1-KO cells compared to WT, while replication was enhanced in cells complemented with TREX1 (Fig 6G). TREX1 does not directly affect IAV polymerase activity, eliminating a trivial explanation for changes in viral replication measured with our reporter virus (Fig S6A). Cells transfected with total nucleic acids were almost completely refractory to viral infection and replication, independent of the presence of TREX1 (Fig 6G), reflecting the potent immunogenicity of the RNA ligands in extracts from infected cells (Fig 6A). Crucially, cells transfected with RNA-depleted samples showed a TREX1-dependent phenotype. Viral replication was restored to moderate levels only in complemented cells (Fig 6G), which express TREX1 above endogenous levels of TREX1 (Fig S3F). These are the same conditions where TREX1 complementation partially suppresses innate sensing of host DNA (Fig 6B, D). Treatment of extracts with RNAse and DNAse restored replication back to untransfected levels. Combined, our results show that TREX1 acts at the top of the DNA-sensing pathway to moderate flux through cGAS and STING and the subsequent activation of a broad-spectrum antiviral program.

## Discussion

We developed TRPPC, a novel competition-based screening approach that offers several advantageous features for gain-of-function screens. First, the pathogen itself drives the screen by modulating gene expression to directly affect its own replicative fitness. The relative fitness of any individual virus, reflected by its abundance in the population, inherently rank-orders the importance of the targeted gene. The sgRNA encoded by the pathogen uniquely identifies the host gene whose expression alters replication, rapidly connecting specific changes in host gene expression to viral replication. Second, TRPPC-mediated transcriptional changes occur only in infected cells after onset of viral gene expression. Thus, TRPPC screens begin in an otherwise naïve cell population and selectively probe the less-studied middle to late stages of replication. Lastly, TRPPC is highly modular and flexible. We already demonstrated its utility for transcriptional activation and inhibition. TRPPC can be easily adapted for targeted screens, epistatic screens where the pathogen encodes sgRNAs concurrently targeting two different host genes, *in vivo* screens in transgenic animals^63^, or new CRISPR technologies as they become available. Moreover, the TRPPC approach is not restricted to influenza virus, or even viruses in general. The strategy is agnostic to the pathogen; TRPPC-style screening can be performed in any system that can encode and deliver an sgRNA. As there is a clear need for the development of new anti-pathogen therapies and the identification of host-based anti-pathogen targets^4^, TRPPC addresses many of our current blind spots for discovering these host dependency factors.

The unbiased screening and focus on the middle to late stages of replication inherent to TRPCC screens have the potential to identify unappreciated biology that affects viral replication. For example, *SLC9C1* was the top-ranked candidate from our screen (Fig 2E, 2G). This was surprising as this gene is not normally expressed in lung cells^64^. SLC9C1 (sNHE) is a sperm-specific Na+/H+ exchanger that controls intracellular pH^65^. We also identified guanine nucleotide–binding protein–coupled estrogen receptor 1 (GPER1) as a target enriched at least 2-fold in two of our screens (Supplemental Table 1). GPER1 is abundantly expressed in reproductive and fetal tissues, and it was previously shown that GPER1 activity suppresses IFN signaling^66^. Neither of these genes are normally expressed in A549 cells, the cells used for our screen^67^. This highlights the benefits of gain-of-function strategies and the ability of TRPPC to survey the full genome, not just those genes that are expressed in any particular cell type. While the mechanism by which SLC9C1 affects IAV replication is unclear, its appearance as a top candidate in all three independent screens suggests a strong impact on viral replication and raises the possibility of previously unappreciated that impact virion production. A potential limitation of the screen and the interpretation of results involves the CRISPRa/i system. The efficiency of transcriptional control is affected by the sgRNA, some of which are more potent than others. Thus, enrichment of any particular sgRNA will be a combination of the effect its target gene has on viral replication, and how potently it alters expression of that target. This might help to explain results from our genome-wide screen. Replicate screens were highly reproducible (Fig 2F, Supplemental Table 1). Yet, within a screen, only one sgRNA was enriched for most targets. We addressed this in part by incorporating sgRNAs designed under different rules into our gene-specific validation experiments, and this approach could be expanded to larger sub-libraries of top candidates for recursive screening. Recurring selection of a target gene, independent of the sgRNA used, would prioritize that target for mechanistic studies.

TREX1 normally functions at the top of the DNA sensing cascade to metabolize DNA in the cytoplasm and prevent inappropriate activation of the innate immune system^37,43^. TREX1 dysfunction can cause chronic autoinflammatory disease^45^. The pro-viral function of TREX1 derives from this same activity. TREX1 degrades immunogenic viral DNAs made during HIV-1 infection, suppressing innate immune activation and increasing viral replication^43,44^. The current work reveals TREX1 also clears host mtDNA during viral infection before it is recognized by innate immune nucleic acid sensors (Fig 5). We and others have shown IAV infection triggers mtDNA leakage (Fig 5A)^53^, and we now show TREX1 regulates the abundance and sensing of mtDNA to promote replication (Fig 5B). Our results suggest that this activity of TREX1 may have a generalizable pro-viral function where infections induce mtDNA stress or release, as is the case for multiple herpesviruses, dengue virus, SARS-CoV-2, chikungunya virus, and Zika virus^54,54,55,68–71^. Activating the orthogonal pathways of RNA- and DNA-sensing amplifies innate immune responses and provides resilience to viral countermeasures. Nonetheless, flaviviruses and alphaviruses encode proteins that counteract this antiviral response by targeting both cGAS and STING for degradation, highlighting the importance of DNA sensing in controlling RNA virus infection^55,69,71,72^.

The proximal cause of mtDNA leakage during IAV infection is not clear. Multiple viral proteins have been implicated in disrupting mitochondria integrity. The polymerase subunit PB2 can localize to the mitochondria via an N-terminal mitochondrial-targeting signal^73^. PB1-F2, produced from an alternative open reading frame in *PB1*, also targets and disrupts the mitochondrial membrane^74^. But, these properties are not conserved in all influenza virus isolates^75,76^, and are absent in some of the primary isolates we used. TREX1 enhanced replication for all the strains we tested, excluding a necessary role for mitochondria-localized PB2 or PB1-F2. The viral pH-activated ion channel M2 has also been suggested to permeabilize the mitochondrial membrane and release mtDNA^53^. But again, we demonstrated a pro-viral effect of TREX1 for IBV and VSV, neither of which encode a direct homolog of IAV M2. This supports the possibilty that the stress of infection such as the unfolded protein response (UPR) or inflammatory cytokine signaling, as opposed to a specific viral protein, is sufficient to induce mtDNA release^77,78^.

Defining the host:pathogen interface is essential to understanding how influenza virus co-opts cellular factors and evades innate antiviral responses. The development of TRPPC screens, where viral fitness selects the highest-confidence candidates, has shown that important regulators of infection like TREX1 remain to be identified. These data highlight the power of pathogen-driven screens in implicating new classes of host factors.

## Experimental Procedures

### Cells

A549 cells (ATCC), Madin-Darby canine kidney (MDCK; ATCC), human embryonic kidney 293T (HEK293T; ATCC), MDCK-SIAT1-TMPRSS2 cells^79^, A549 MAVS knockout cells^80^, and A549 STING knockout cells^72^ were grown at 37°C and 5% CO_2_ maintained in Dulbecco’s modified Eagle’s medium (DMEM) supplemented with 10% fetal bovine serum (FBS) and 1% penicillin-streptomycin. A549 TREX1 knockout cells were generated by Synthego and used to produce clonal knockout lines by limiting dilution. Clonal A549 TREX1-KO cells were screened via western blot and validated by sequencing the expect edit site (Fig S3E-F). All knockout cell lines were paired with parental WT controls from the same source. A549-Cre-reporter CRISPR-SAM clone #15, referred to here as A549-CRISPRa cells, is a clonal cell line expressing dCas9-VP64 and MS2-p65-HSF1^19^. Cultures were routinely tested for mycoplasma using MycoAlert (Lonza).

### Plasmids

The TRPPCa reporter plasmid p9X-NL1.2 was based on pNL1.2[*NlucP*] (Promega) that encoded *Nanoluciferase* fused to a C-terminal PEST sequence from mouse ornithine decarboxylase. A 39 bp fragment compromisin a minimal the CMV immediate early promoter was placed upstream of NLuc and preceded by 9 copies of a 20 bp protospacers sequence ATCTAGATACGACTCACTAT along with *NGG* PAMs suitable for dCas9 recognition^81^.

Expression plasmids for RNP components (PB2, PB1, PA, and NP) used in TRPPCa reporter and polymerase activity were derived from A/WSN/33 (H1N1; WSN)^82^.

Genomic segments for rescue of A/PuertoRico/8/34 (H1N1; PR8) viruses were expressed from plasmids pHW190-PB2 to pHW198-NS (a kind gift from S. Schultz-Cherry). Split-NS was generated as previously-described^28^. pHW-TRPPC-NS plasmids were generated by placing mir124 into the artificial intron as before^28^ and inserting sgRNA2.0 downstream^26^ (Fig S7). sgRNA2.0 contains two hairpins recognized by MS2 coat protein. Specific sgRNA sequences were cloned via inverse PCR or ligation of self-annealing oligo pairs into compatible digested vector (Supplemental Table 2). The TRPPC-NS construct used in reporter assays recognizes the target sequence ATCTAGATACGACTCACTAT^81^.

RNP components for A/California/07/2009 (H1N1; CA07) and B/Brisbane/60/2008 (B/Bris) used in TRPPCa reporter assays were expressed using bidirectional pHW plasmids^15,83,84^. Split-NS and TRPPC-NS for both CA07 and B/Bris were cloned as described for PR8.

TRPPCi reporter plasmids were cloned from previously described CRISPRi reagents^29^. The reporter includes a Nluc-PEST gene driven by a minimal CMV promoter with a Gal4 upstream activation site and contains the protospacer sequence TACCTCATCAGGAACATGT followed by a PAM TGG. Expression constructs include a VP16-Gal4 transactivator, a dCas9-KRAB-MeCP2 repressor, and a TRPPCi-NS plasmid expressing a targeting or control sgRNA sequence.

pHAGE2–EF1aInt–TMPRSS2–IRES–mCherry-W was previously describe^79^.

Human TREX1 sequences were derived from plasmid GFP-TREX1 (Addgene 27219) and TREX1-D18N (Addgene 27220).

Lentivirus packaging plasmids included psPAX2 (Addgene 12260), pCMV-VSV-G (Addgene 8454), pLenti dCAS-VP64_Blast (Addgene 61524), pLenti MS2-P65-HSF1_Hygro (Addgene 61426), and pLX304 (dCas9-VP64, MS2-p65-HSF1). The UCOE-SFFV backbone (Addgene 122205) was used to create pSFFV-TREX1-V5-2A-BSD and pSSFV-TREX1-D18N-V5-2A-BSD.

Plasmids for Sleeping Beauty transposition included pCMV(CAT)T7-SB100 (Addgene 34879) and an ISG54 promoter driving Nluc-2A-GFP cloned into the pSBbi-BB backbone (Addgene 60521).

### Stable cell lines

VSV-G pseudotyped lentiviruses were prepared by co-transfecting 293T cells with psPAX2, pCMV-VSV-G, and either pLenti dCAS-VP64_Blast, pLenti MS2-P65-HSF1_Hygro, pLX304, pSFFV-TREX1-V5, or pSSFV-TREX1-D18N-V5.

Polyclonal 293T-CRISPRa cells were generated by transducing cells with lentiviruses expressing dCas9-VP64 and MS2-p65-HSF1. CRISPRa cells were maintained in media containing 4 μg/ml blasticidin and 150 μg/ml hygromycin. A549 cells overexpressing or complemented with V5-tagged TREX1 or TREX1-D18N were generated by transducing cells and selecting with 6 μg/ml blasticidin. ISRE reporter cells lines were generated using the Sleeping Beauty transposase system^85^ to integrate an ISRE promoter driving NanoLuc-2A-GFP and selected with 6μg/ml blasticidin.

All plasmids were verified by sequencing.

### Viruses and infections

Influenza viruses were derived from A/PR8/34 (H1N1; PR8), A/WSN/33 (H1N1; WSN), A/green-winged teal/Ohio/175/1986 (H2N1; S009), A/California/04/2009 (H1N1; CA04), and B/Brisbane/60/2008 (B-Victoria lineage; B/Bris). Nanoluciferase-expressing reporter viruses express PA-2A-NLuc from the PA segment^15,86,87^. These are collectively referred to as PASTN. VSV-GFP was previously reported^88^.

Cell culture infections were performed in triplicate at 37°C (influenza A virus, VSV) or 33°C (influenza B virus). Cells were inoculated at the indicate MOI with virus diluted in OptiVGM (OptiMEM, 1% penicillin/streptomycin, and 0.2% BSA). For trypsin-dependent influenza viruses, media was further supplemented with 0.5-2 μg/ml TPCK-treated trypsin and 100 μg/ml CaCl_2_.

### Generation of recombinant influenza virus

Individual influenza viruses in the PR8 background were rescued using plasmid-based reverse genetics^89^. Briefly, bidirectional pHW plasmids for each viral genome segment were co-transfected wit pHAGE2–EF1aInt–TMPRSS2–IRES–mCherry-W into 293T cells using jetPRIME (PolyPlus). 24 hr post-transfection, media was removed and cultures were overlaid with MDCK-SIAT1-TMPRSS2 cells in OptiVGM supplemented with 100 μg/ml CaCl_2_. Rescue supernatants were harvested 48 hours later. Individual viruses were plaque-purified by serial dilution on MDCK cells with agarose overlays. Plaque-purified recombinant viruses were verified by RT-PCR and Sanger sequencing. Higher titer stocks were generated by infecting plates of MDCK-SIAT1-TMPRSS2 cells at low MOI for 24-48 hours.

### Plaque assays and multicycle replication

Viral titers were measured by plaque assay on MDCK cells (influenza virus) or A549 cells (VSV) following prior approaches where cells are overlaid with medium containing 1.2% Avicel (catalog number RC581; FMC BioPolymer)^90^.

For multicycle replication kinetics of VSV-GFP and influenza viruses, cell lines were infected in triplicate at the indicated MOIs. Aliquots were removed throughout the infection and titers were determined by plaque assays. For bioluminescent reporter viruses, viral titers were determined by infecting MDCK cells and measuring reporter activity using the Nano-Glo luciferase assay (Promega)^86,91^. Viral replication in the presence of a STING agonist was performed by pre-treating cells with diABZI (28054; Cayman Chemical) dissolved in DMSO for 4 hours prior to infection.

### TRPPC reporter assays

293T-CRISPRa or A549-CRISPRa cells were simultaneously seeded and transfected with Transit2020 (Mirus) in triplicate. For transfection-only assays, cells were transfected with the reporter plasmid p9X-NL1.2, pHW198-NS or variant, and expression vectors for viral NP and polymerase proteins where indicated. Forty-eight hours post-transfection, cells were lysed in coIP buffer (50 mM Tris, pH 7.4, 150 mM NaCl, 0.5% NP-40). Bioluminescence was measured by NanoGlo luciferase assays (Promega). For infection-based assays, cells were transfected only with the reporter plasmid p9X-NL1.2 and subsequently infected 24 hours later with the indicated viruses. Bioluminescence was measured at various timepoints post-infection. For TRPPC-inhibition (TRPPCi), 293T cells were transfected with expression constructs for VP16-Gal4, dCas9-KRAB-MeCP2, viral NP and polymerase proteins, pHW198-NS or variant, and the CRISPRi reporter plasmid. Bioluminescence was assayed 96h post-transfection.

### Reverse transcription qPCR

Total RNA was extracted from cells using Trizol and mRNAs were reverse transcribed using an oligo-dT primer and MMLV reverse transcriptase. cDNA was combined with iTaq SYBR master mix (Biorad) along with gene-specific primers listed in Supplemental Table 3 and qPCR was performed in technical triplicate on a StepOnePlus (Stratagene). Ct values of target genes were normalized to β-actin and relative gene expression levels between conditions were calculated via the ΔΔCt method. Expression levels are plotted as fold-change over a baseline condition set to 1. Values are the mean of 3 biological replicates.

### Generation of genome-wide TRPPC virus library

The human SAM CRISPRa sgRNA library (Addgene #1000000078) was cloned into the pHW-TRPPC-NS rescue plasmid backbone for PR8 (Fig S2A, S7). sgRNAs were PCR-amplified from the SAM library in 24 low-cycle reactions using primers that append restriction sites compatible with the pHW-TRPPC-NS cloning vector. PCR products were pooled and concentrated using DNA Clean and Concentrate columns (Zymo). Insert (sgRNAs) and vector (pHW-TRPPC-NS) were digested with restriction enzymes to generate compatible sticky ends and gel-purified. Fragments were ligated in 20 separate reactions, transformed into Mach1 competent E. coli (Thermo), plated on LB agar supplemented with 100 μg/ml ampicillin, and grown overnight at 37°C. The number of transformants was ∼1,118,500. Colonies were scraped off plates into 1 L of liquid LB containing ampicillin and incubated for 3 hours at 37°C. DNA was purified using a MaxiPrep Kit (Macherey-Nagel). Plasmid stocks were deep-sequenced to ensure complete representation.

pHW-TRPPC-NS plasmids undergo low-level recombination in bacteria due to direct repeats in the split *NS* segment, recreating WT *NS*. To avoid this contaminant and generate virus libraries exclusively containing the TRPPC NS segment, plasmid stocks were linearized with PvuI and size-selected on agarose gels to ensure removal of smaller recombined NS plasmids (Fig. S2A). Linearized TRPPC-NS plasmid was used with the other seven PR8 rescue plasmids pHW190-PB2 to pHW197-M in 90 individual transfection that were subsequently pooled and titered. Samples from this primary supernatant were DNAse-treated and analyzed by deep-sequencing to determine library size and composition before starting TRPPC screens (Fig S2B). 69,276 unique TRPPC viruses were detected. Population diversity was measured using Shannon’s diversity (H’)^92^, as follows:

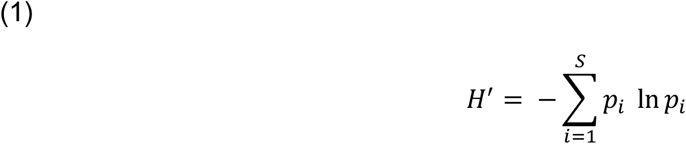

### TRPPC-NS stability assessment

A549 cells were inoculated with TRPPC virus libraries at an MOI of 0.05. Supernatants were harvested 48 hours later from which aliquots were used to re-initiate another round of infections. This was repeated for a total of 4 passages. Viral RNA was then extracted from all samples with Trizol and reverse transcribed SuperScriptIV (ThermorFisher) with a primer specific for NS vRNA. cDNA was PCR amplified and products were imaged on agarose gels next to amplicons from WT and Split-NS segments as size comparisons.

### TRPPC miniscreen and competition assays

34 unique TRPPC viruses each containing a different sgRNA were individually rescued and titered. This included a non-targeting virus and viruses targeting 11 different genes, each with three distinct sgRNAs. These viruses were pooled in equivalent amounts based on plaque forming units. The competitive screen was performed in duplicate by inoculating CRISPRa-A549 cells at an MOI of 0.05. Supernatants were harvested 48 hpi, titered by plaque assay, and used to initiate a subsequent round of competition. This process was repeated for a total of 4 rounds. Complementary DNA (cDNA) was generated from RNA samples by reverse transcription using SuperScript IV (ThermoFisher) and NS-specific primers that appended a 10 nt (NNNNNNNNNN) unique molecular identifiers (UMIs) to each TRPPC-NS segment. cDNA was amplified by PCR with primers that appended sequencing adapters and indices. Amplicons were pooled and sequenced on an Illumina MiSeq, NextSeq 500, or NovaSeq 6000. Replicate reads were collapsed into single UMIs and counted before data analysis (analysis pipeline detailed below).

For the miniscreen, sequencing data were used to quantify the abundance of each sgRNA in the pool before and after competition. As plaque forming units do not directly measure genome abundance, the frequency of each viral genome varied from perfect equivalence in the starting population. Therefore, the frequency of each virus in the starting pool was normalized to 1, and their relative enrichment or depletion was examined over the serial passages.

For head-to-head-competition with TREX1-targeting TRPPC viruses, equal plaque forming units of 3 TREX1 viruses each encoding a different sgRNA, 2 non-targeting viruses, and an MX1-targeting virus were pooled. A549-CRISPRa cells were inoculated with pooled virus at an MOI of 0.05. Supernatants were harvested 48 hpi. Amplicons of the genomic NS segments from input and output population were prepared as above and sequence. The frequency was determined for each virus before and after competition. Competition experiments was performed in biological duplicate.

### Genome-wide TRPPC screens

240 million CRISPRa-A549 cells were inoculated at an MOI of 0.05 with the genome-wide TRPPC-NS virus library for a theoretical library coverage of 170x for each sgRNA. Supernatants were collected 48 hpi, pooled, clarified, titered by plaque assay, and vRNA was extracted with Trizol for subsequent deep sequencing. Supernatants were then used to initiate serial rounds of infection under the same conditions, for 5 total rounds. To avoid technical bottlenecks in preparing a large library, UMI-tagged amplicons were created as above in 6 separate reverse-transcription reactions that were pooled and amplified in 12 separate PCRs. Amplicons were pooled and sequenced. Beginning with the same starting library, the entire screen was performed in triplicate.

### Deep sequencing data analysis of viral genomes in TRPPC screens

Next-generation sequencing data was processed using the following pipeline. Data quality control filtering was done with FastQC v.0.11.5. Reads were merged with BBMerge from the BBMap (v38.49) suite^93,94^. Adapters were trimmed with BBDuk, also from the BBMap (v38.49) suite. Reads were aligned with bowtie2 (v2.3.5.1) against a custom index containing the sgRNA sequences expected to be present in each experiment^95^. PCR duplicates were removed by collapsing reads based on UMIs to yield a unique number of viruses mapping to each sgRNA. Python (v3.7.3) was used to automate processing. For the genome-wide screen, all three replicates were used for to identify gene targets enriched over the course of selection using MAGeCK (v0.4)^96^. GO analysis was performed with ShinyGO^97^.

### Immunofluorescence

WT, TREX1-KO, or complemented A549 cells were seeded on glass coverslips and infected the next day with PR8 (MOI = 1). At the indicated timepoints, coverslips were washed with PBS and fixed with 4% paraformaldehyde. Cells were permeabilized with 0.1% Triton X100, blocked with 1% BSA in PBS, incubated with primary antibodies anti-RNP (1:1000, BEI Resources Repository NR-3133) and anti-dsDNA (1:100, Santa Cruz Biotechnology sc-58749), and then secondary antibodies donkey anti-goat 488 AlexaFluor (Invitrogen) and chicken anti-mouse 594 AlexaFluor (Invitrogen). Images were acquired on Zeiss confocal fluorescence microscope.

### Western blotting

Cell were lysed in CoIP buffer containing protease inhibitor cocktail (Roche) and clarified by centrifugation. Samples were separated via denaturing polyacrylamide gel electrophoresis and transferred to polyvinylidene fluoride membranes prior to blocking with 5% milk and incubation with antibodies. Primary antibodies used were: anti-TREX1 (1:500, Proteintech), anti-RNP (1:1000, BEI Resources Repository, NR-3133), anti-V5-HRP (1:10000, Sigma), and anti-tubulin (1:5000, Sigma). Secondary antibodies were HRP-conjugated and chemiluminescent images were acquired with an Odyssey Fc Imager equipped with Image Studio (LI-COR). Experiments were performed in triplicate with representative images selected.

### Polymerase activity assays

Activity assays were performed as described^98^, where HEK293T cells were simultaneously seeded and transfected in technical triplicate with plasmids expressing RNP components from WSN (PB2, PB1, PA, NP), a firefly luciferase reporter in the context of the vNA segment, an SV40-driven Renilla luciferase control, and either GFP-TREX1 or GFP alone. Cells were lysed 48h post-transfection in Renilla lysis buffer (Promega) and bioluminescence was measured for both firefly and Renilla luciferases for each sample. Firefly was normalized to background Renilla values.

### Mitochondrial DNA detection assay

Detection of mtDNA in cytosolic extracts was performed as described previously^53^. Briefly, cells were harvested and gently pelleted by centrifugation. Pellets were resuspended in buffer comprised of 150 mM NaCl, 50mM HEPES pH 7.4, and 20 μg/mL digitonin. Samples were gently nutated for 10 minutes followed by centrifugation at 1000 x g for 10 minutes. Supernatants were transferred to fresh tubes and centrifuged at 17000g for 10 minutes. DNA in these cytosolic fractions was concentrated with a DNA clean and concentrator kit (Zymo). Matched samples containing total mtDNA were isolated from whole cell extracts using DNeasy Blood and Tissue kits (Qiagen). qPCR was performed on both the cytosolic fraction and whole cell extract for each experimental condition using mtDNA specific primers: Fwd 5’-CCTAGGGATAACAGCGCAAT-3’, Rev 5’-TAGAAGAGCGATGGTGAGAG-3.’ Relative cytosolic mtDNA levels were normalized to whole cell extracts and the value in mock-infected WT cells was set to 1.

### ISRE reporter assays

ISRE reporter cells lines based on WT or TREX1-KO A549 cells were used to measure innate immune activation. Cells were either simultaneously seeded and transfected with various nucleic acids using Transit-X2 (Mirus), seeded and treated with IFN? 24 h later, or seeded and inoculated 24 h later with with influenza virus. Cytosolic DNA from infected A549 cells was acquired by preparing cytosolic extracts with digitonin as above, extracting DNA with phenol/chloroform/isoamyl alcohol, followed by precipitation and resuspension. Human DNA from whole cell extracts of A549 cells was purified using the DNeasy Blood and Tissue kit (Qiagen). RNAseA (Thermo) and DNAseI (Promega) were used to treat nucleic acid extracts where indicated. Poly(I:C) and salmon sperm DNA ligands were acquired from Sigma. Followin treatment, cells were lysed in CoIP buffer and bioluminescence was measured via NanoGlo luciferase assays (Promega). ISRE induction values were normalized to untreated cells for each cell line.

### RNA sequencing

Total RNA was isolated from TREX1-KO or complemented A549 cells that were either mock- or PR8-infected (MOI = 0.5) for 24h. Samples were DNAse-treated and sent for library preparation and mRNA-seq by NUcore (Northwestern) on a HiSeq 4000 (Illumina) with 50 bp single-end reads. Sequencing was performed in biological triplicate. Python anaconda 3.6 was used to trim sequences with Trim Galore (v0.4.4)^99^, align with STAR (v2.5.3a)^100^, and count with HTSeq (v0.9.1)^101^. Differential gene analysis was performed in R (v4.1.1) running DESeq2 (v1.34.0)^102^. Analysis of ISGs utilized a previously reported extended gene set^21^. GO analysis was performed with ShinyGO^97^.

### Data Availability

All sequencing files have been deposited as BioProject PRJNA930886 for the screen and PRJNA930919 for RNA-seq. Source data are provided in this paper with underlying raw data and uncropped blots in Supplemental Table 4 and 5.

### Statistical Analyses

Statistical analyses was performed in GraphPad Prism (v9.4.1). Pairwise comparisons were performed using an unpaired two-way Student’s T-test. Multiple comparison were made via ANOVA followed by post-hoc analysis using Šídák’s, Tukey’s or Dunnett’s correction as indicated. Correlation between screens was measured with Spearman’s rank correlation (ρ). All experiments were performed in technical triplicate with at least three biological replicates, with the exception of the competitions and screens that had 2-3 biological replicates. Figures were assembled in Adobe Illustrator (27.0.1).

## Acknowledgements

We thank Jesse Bloom, Alex Chavez, Nicholas Heaton, Craig McCormick, Sara Sawyer, Stacey Schultz-Cherry for generously sharing reagents. We acknowledge Eva Berlot and members of the Mehle lab, especially Mitch Ledwith, for critical input. CRK was an Open Philanthropy Fellow of the Life Sciences Research Foundation. AM is a Burroughs Wellcome Fund Investigator in the Pathogenesis of Infectious Disease and an H.I. Romnes Faculty Fellow funded by the Wisconsin Alumni Research Foundation (USA). This work was supported by National Institutes of Health grants AI125897 and AI164690 to AM, AI007414 and GM140935 to GAS, and GM007215 to KAA.

