## Supplemental Figures 1-7 for "Pathogen-driven CRISPR screens identify TREX1 as a regulator of DNA self-sensing during influenza virus infection"

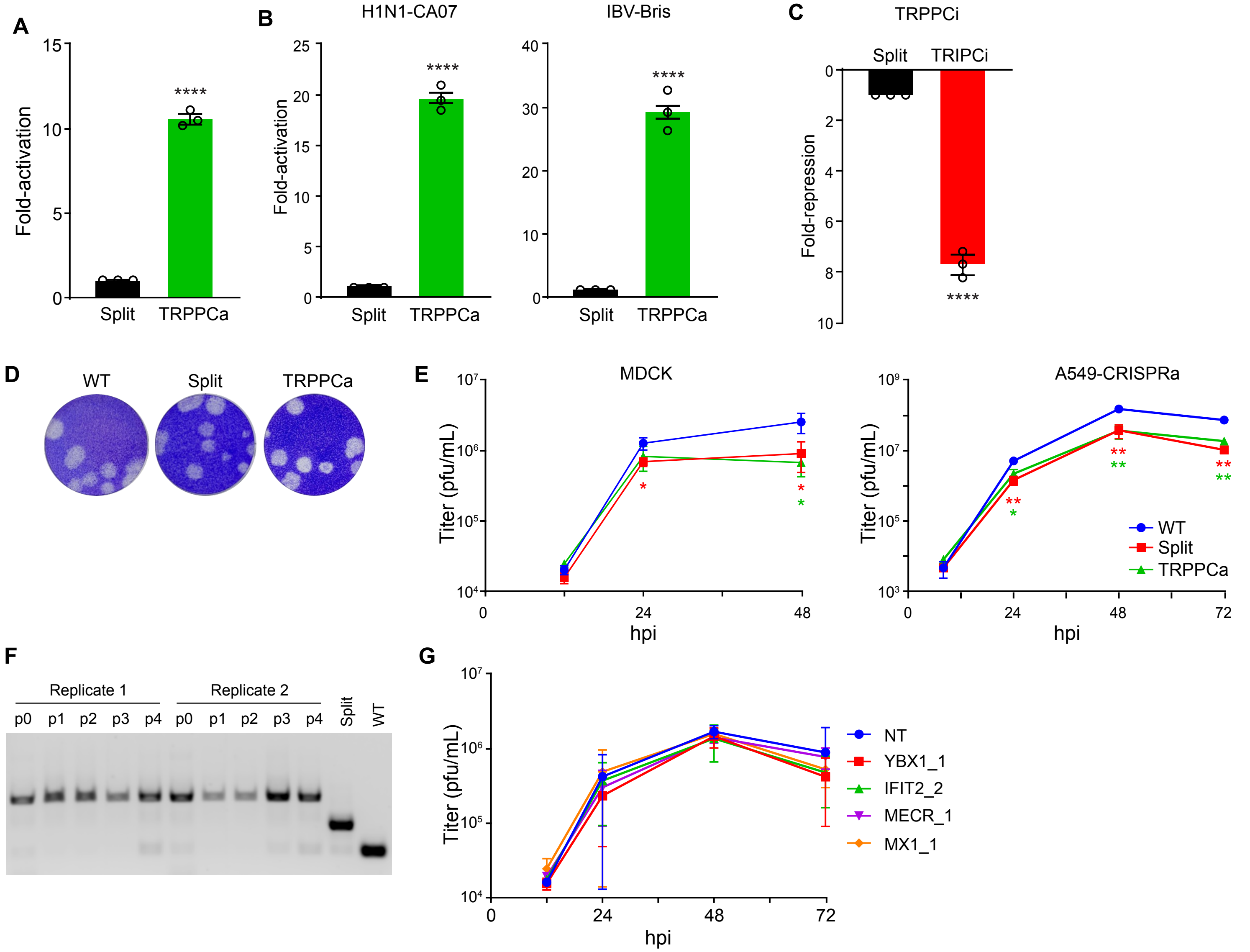
A
Fold-activation
Split
TRPPCa
****
B
H1N1-CA07
IBV-Bris
Fold-activation
Split
TRPPCa
****
C
TRPPCi
Split
TRIPCi
Fold-repression
****
D
WT
Split
TRPPCa
E
MDCK
A549-CRISPRa
Titer (pfu/mL)
hpi
WT
Split
TRPPCa
F
Replicate 1
Replicate 2
p0
p1
p2
p3
p4
Split
WT
G
Titer (pfu/mL)
hpi
NT
YBX1_1
IFIT2_2
MECR_1
MX1_1

**Figure S1 Extended validation of TRPPC system.**

**A,** TRPPCa in A549-CRISPRa cells of a luciferase reporter targeted by sgRNA expressed from transfected *NS* in the presence of the viral replication machinery. *Split NS* that lacks an sgRNA is included as a control.

**B,** TRPPCa functions with *NS* from primary isolates of IAV and IBV. Activation of a luciferase reporter targeted by sgRNA expressed from transfected CA07 or IBV *TRPPC-NS* in the presence of matched viral polymerase and NP.

**C,** TRPPC-inhibition (TRPPCi) suppresses gene expression. PR8 *TRPPC-NS* suppressed reporter

**D,** Example plaque morphologies for WT and engineered viruses.

gene expression when transfected into cells with the viral polymerase, NP, and dCas9-KRAB.

**E,** TRPPC viruses replicate similar to WT in multiple cell lines. Multicycle replication kinetics of WT, split-NS, or TRPPCa-NS with a non-targeting sgRNA in MDCK and A549-CRISPRa cells (MOI = 0.01).

**F,** Engineered *NS* segment stability was measured over serial passages by assessing amplicon sizes by RT-PCR.

**G,** TRPPC targeting does not affect replication in cells lacking the CRISPRa machinery. Multicycle replication of TRPPC viruses targeting specified host genes in WT A549 cells inoculated at MOI = 0.01.

Data are shown as grand mean of 3 replicates ± SEM (C, D, F) mean ± s.d. (E, G). Unpaired T tests (C, D, F) or one-way ANOVA with post-hoc Dunnett's tests (E, G) were performed (*p<0.05, **p<0.01,  ****p<0.0001).

A

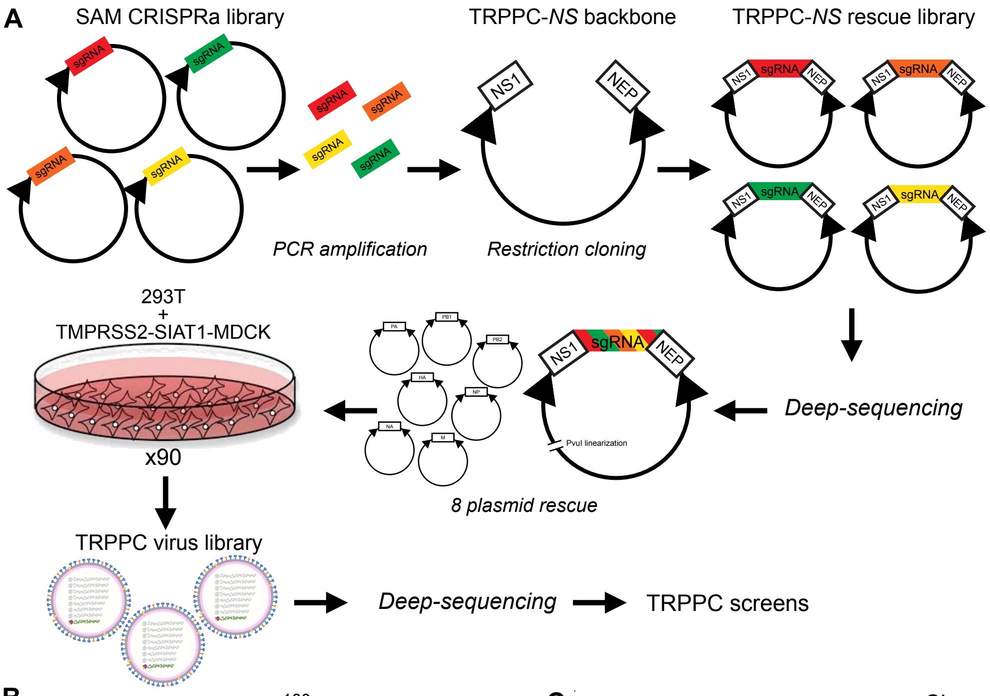


B

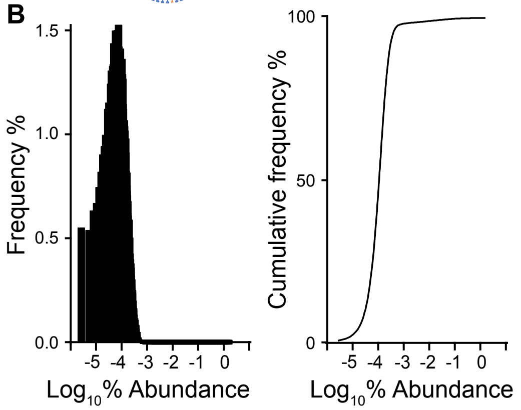


C

### Virus

Shannon Diversity (H′)

plasmid
10.90131

| p0 | p1 | p2 | p3 | p4 | p5 |
|---|---|---|---|---|---|
| 6.506059 | 5.682367 | 5.282353 | 5.192416 | 5.068081 | 4.850333 |
| 6.506059 | 5.678302 | 5.305056 | 5.20663 | 5.122207 | 4.955211 |
| 6.506059 | 5.646149 | 5.202897 | 5.077303 | 4.987117 | 4.769359 |

richness

69179

| p0 | p1 | p2 | p3 | p4 | p5 |
|---|---|---|---|---|---|
| 69276 | 3742 | 1993 | 1768 | 1652 | 1565 |
| 69276 | 3656 | 1989 | 1750 | 1629 | 1555 |
| 69276 | 3512 | 1880 | 1661 | 1536 | 1419 |

D

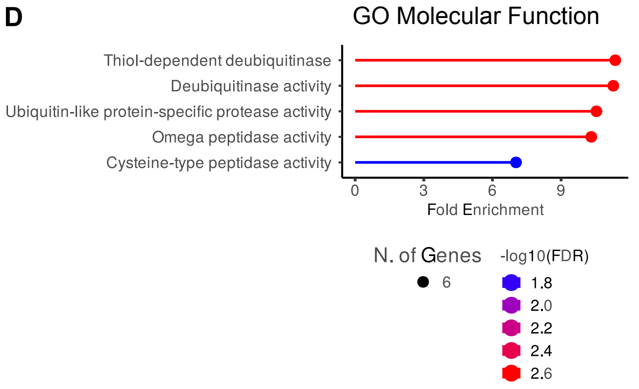


E

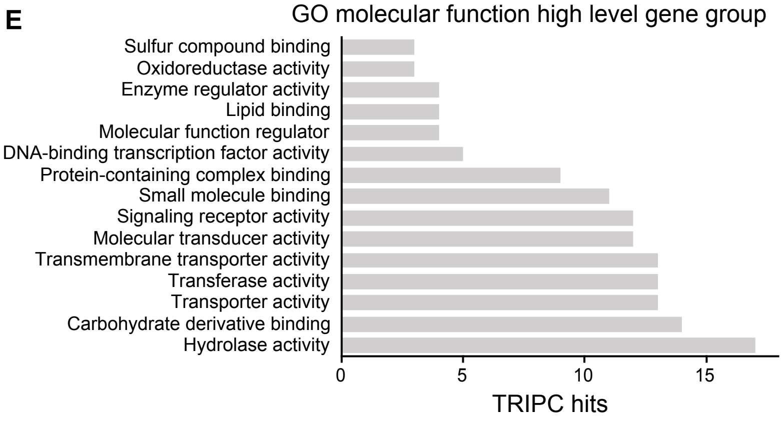

**Figure S2 Characterization of TRPPC library and gene enrichment analysis.**

**A,** Experimental workflow for the creation of a genome-wide TRPPC virus library.

**B,** Distribution histogram and cumulative frequency plot of each member in the TRPPC virus library stock prior to selection.

**C,** Shannon's diversity indices (H') and richness of the viral populations at each passage during the 3 TRPPC screens.

**D,** GO analysis highlighting the molecular function pathways enriched among the top 100 selected genes based on MAGeCK scores.

**E,** Groupings of high-level gene functions conferred by the top 100 genes.

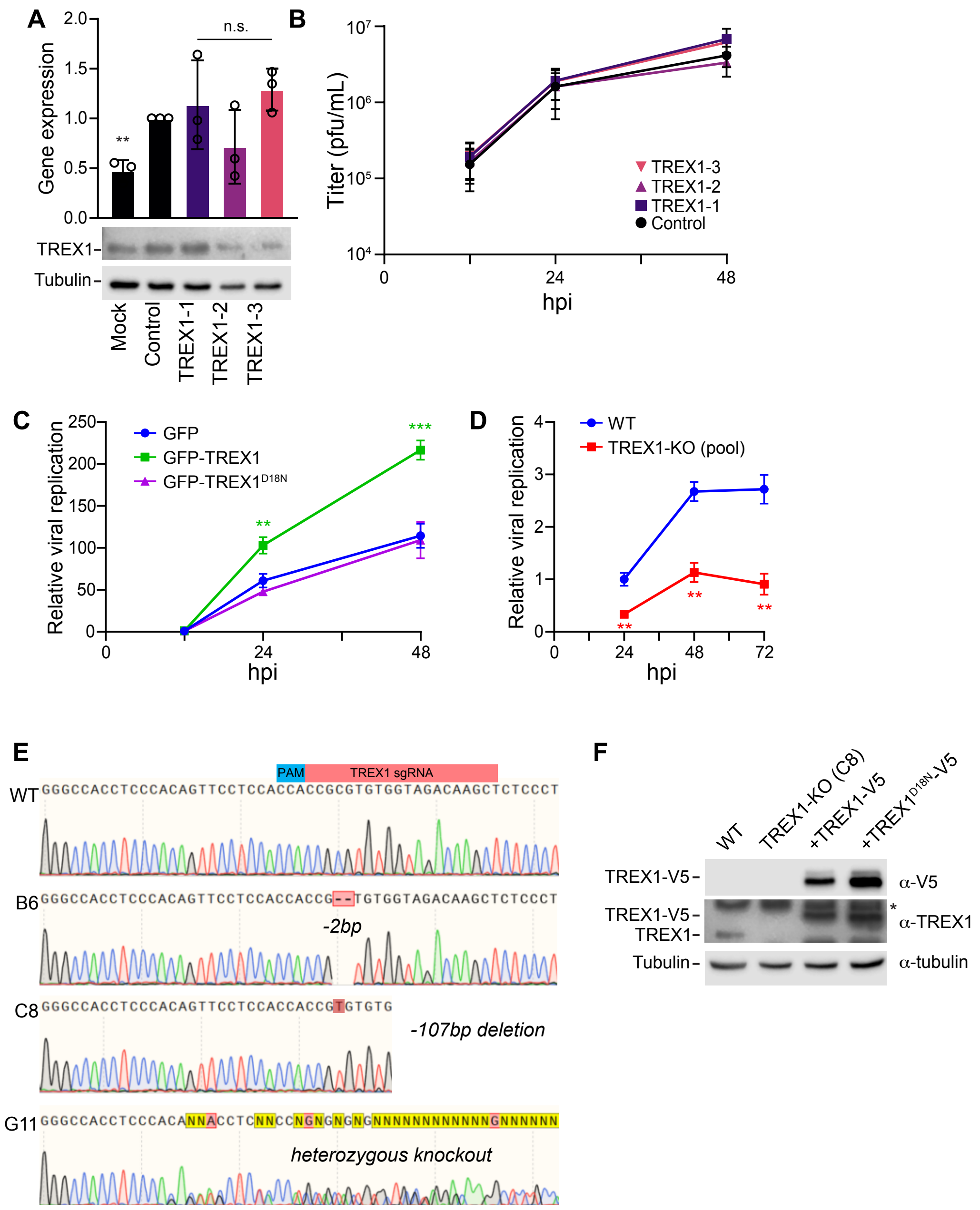
A
Gene expression
2.0
1.5
1.0
0.5
0.0
**
n.s.
TREX1-
Tubulin-
Mock
Control
TREX1-1
TREX1-2
TREX1-3
B
Titer (pfu/mL)
$10^7$
$10^6$
$10^5$
$10^4$
0
24
48
hpi
TREX1-3
TREX1-2
TREX1-1
Control
C
Relative viral replication
250
200
150
100
50
0
0
24
48
hpi
GFP
GFP-TREX1
GFP-TREX1$^{D18N}$
**
***
D
Relative viral replication
4
3
2
1
0
0
24
48
72
hpi
WT
TREX1-KO (pool)
**
**
**
E
PAM
TREX1 sgRNA
WT GGGCCACCTCCCACAGTTCCTCCACCACCGCGTGTGGTAGACAAGCTCTCCCT
B6 GGGCCACCTCCCACAGTTCCTCCACCACCG--TGTGGTAGACAAGCTCTCCCT
-2bp
C8 GGGCCACCTCCCACAGTTCCTCCACCACCGTGTGTG
-107bp deletion
G11
heterozygous knockout
F
WT
TREX1-KO (C8)
+TREX1-V5
+TREX1$^{D18N}$-V5
TREX1-V5-
α-V5
TREX1-V5-
TREX1-
*
α-TREX1
Tubulin-
α-tubulin

**Figure S3 Further validation of TREX1 as a proviral host factor**.

**A,** TREX1 expression was measured in WT A549 inoculated with *TREX1*-targeting or non-targetting control TRPPC viruses (MOI = 1) by RT-qPCR (top) and western blot (bottom).

**B,** Multicycle replication of *TREX1*- or non-targeting TRPPC viruses in WT A549 cells (MOI = 0.01). Titers determined by plaque assay.

**C,** Multicycle replication of a WSN influenza A reporter virus (MOI = 0.05) in A549 cells transfected with GFP-tagged TREX1, $TREX1^{D18N}$ or a GFP-alone control.

**D,** Multicycle replication of a WSN influenza A reporter virus (MOI = 0.05) in A549 cells expressing GFP-tagged TREX1, $TREX1^{D18N}$ or a GFP-alone control.

**E,** *TREX1* genotype of knockout cells. Sanger sequencing traces display CRISPR-Cas9 editing at the *TREX1* locus for 3 selected knockout clones. Edits compared to the WT genome are shown for 2 homozygous (B6, C8) and 1 heterozygous (G11) clones.

**F,** TREX1 knockout was confirmed by western blotting lysates from WT, clonal KO, and complemented A549 cell lines. Endogenous TREX1 and recombinant TREX1-V5-2A are indicated. * = non-specific bands. These cells lines are used throughout Fig 3-6.

Data are shown as grand mean of 3 replicates ± SEM (C-D) or mean ± s.d. (A-B). Unpaired T tests (D) or one-way ANOVA with post-hoc Dunnett's tests (A, C) were performed (*p<0.05, **p<0.01, ***p<0.001, ****p<0.0001, ns = not significant).

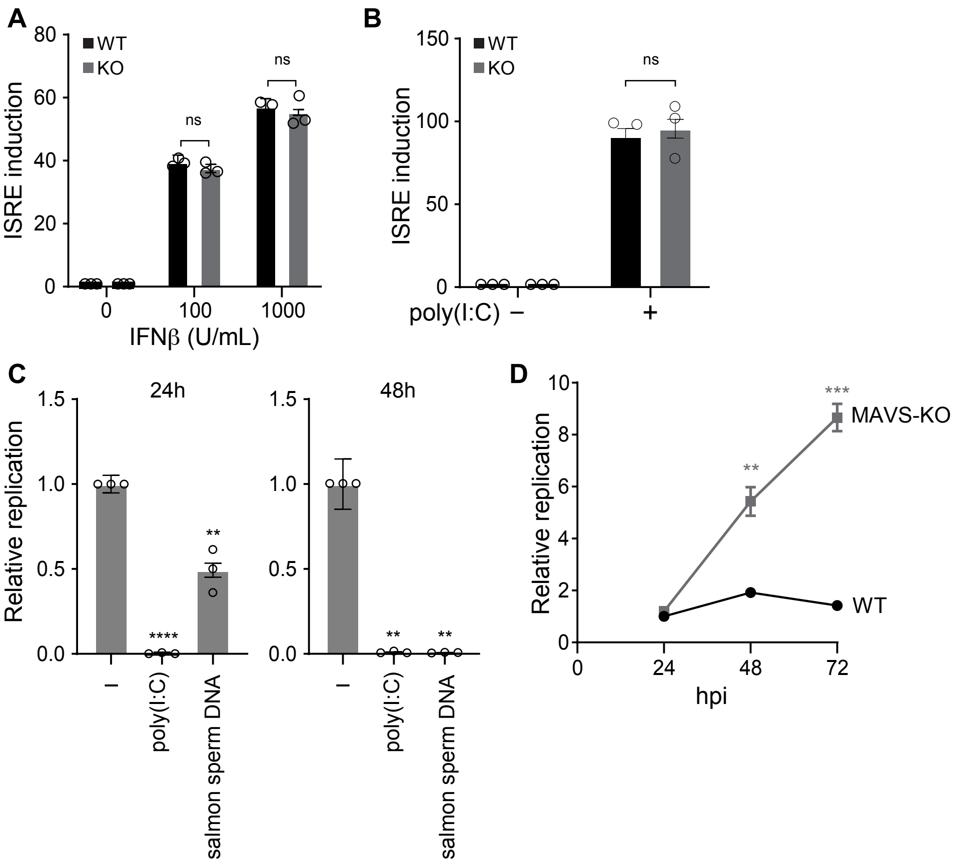
A
WT
KO
ns
ns
ISRE induction
0
20
40
60
80
0
100
1000
IFNβ (U/mL)
B
WT
KO
ns
ISRE induction
0
50
100
150
poly(I:C) −
+
C
24h
48h
Relative replication
0.0
0.5
1.0
1.5
****
**
**
**
−
poly(I:C)
salmon sperm DNA
D
Relative replication
0
2
4
6
8
10
***
**
MAVS-KO
WT
0
24
48
72
hpi

**Figure S4 Reporter cell line validation and IAV replication in MAVS-knockout.**

**A-B,** IFN signaling and RNA sensing remain intact in TREX1-KO cells. ISRE induction in WT and TREX1-KO reporter cells treated with a)IFNβ or b) transfected with poly(I:C). ISRE activation is normalized to untreated and mock-transfected cells, respectively.

**C,** Sensing of foreign nucleic acids blocks IAV replication. Replication of influenza A virus (MOI = 0.05) in WT A549 cells treated with the indicated nucleic acid ligands.

**D,** Infection in cells lacking RNA sensing for comparison. Multicycle replication of influenza A virus (MOI = 0.05) in WT and MAVS-KO A549 cells.

Data are shown as grand mean of 3 replicates ± SEM. Significance was tested with a two-way ANOVA with Šídák's mulitple comparisons (A-B) or a one-way ANOVA with post-hoc Dunnett’s test (**$p<0.01$, ***$p<0.001$, ****$p<0.0001$, ns = not significant).

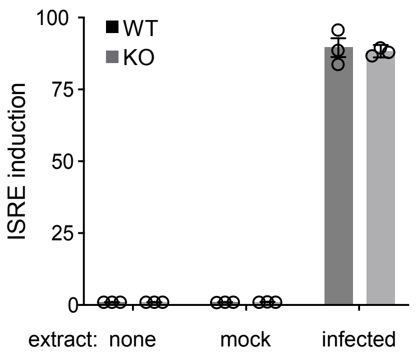
WT
KO
ISRE induction
100
75
50
25
0
extract:
none
mock
infected

**Figure S5 ISRE induction by untreated cytosolic extracts.** ISRE induction was measured in WT and TREX1-KO reporter cells transfected with untreated cytosolic extracts derived from mock or infected cells. Bioluminescence values are normalized to untransfected cells.

Data are shown as grand mean of 3 replicates ± SEM.

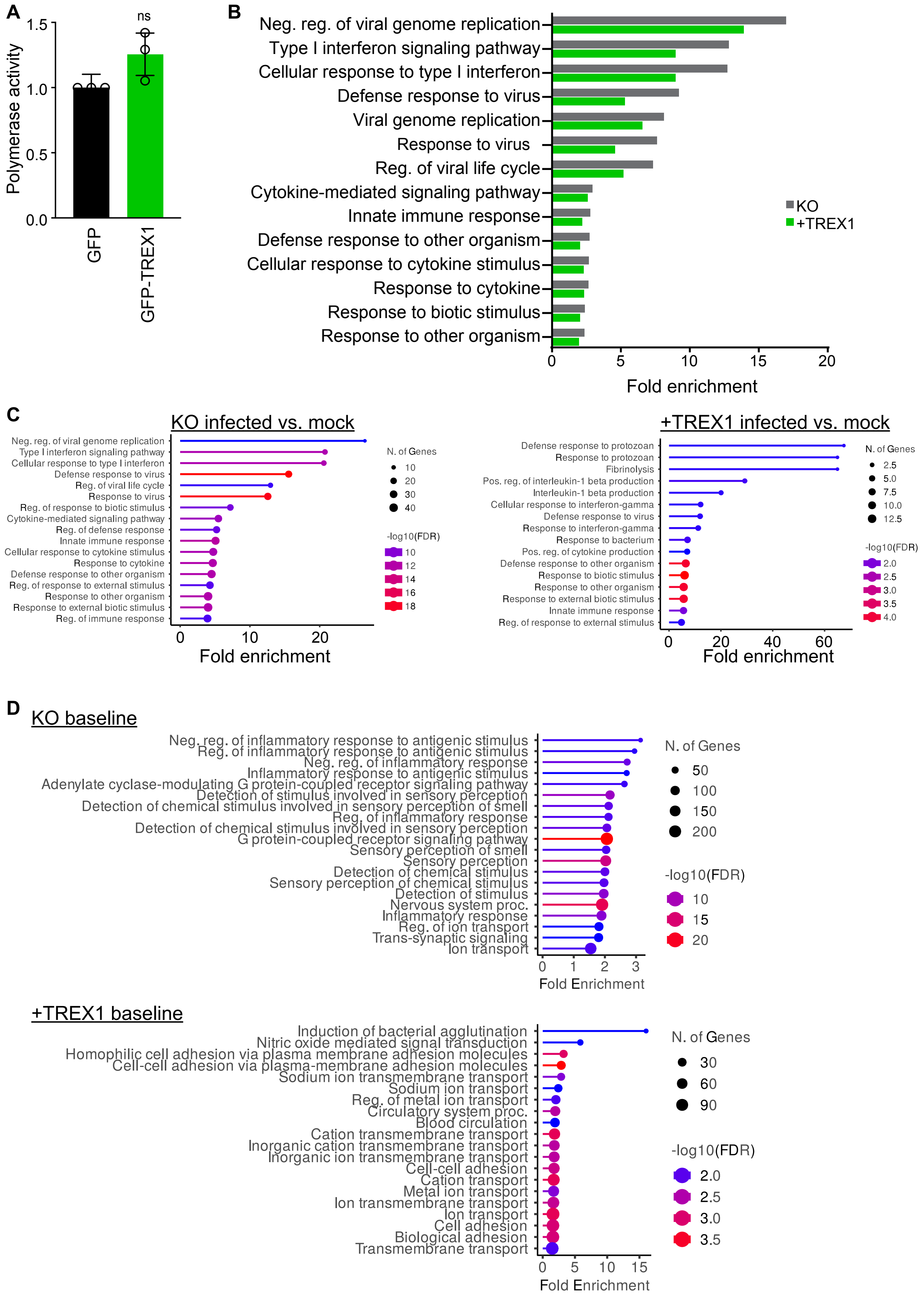
A
ns
Polymerase activity
1.5
1.0
0.5
0.0
GFP
GFP-TREX1
B
Neg. reg. of viral genome replication
Type I interferon signaling pathway
Cellular response to type I interferon
Defense response to virus
Viral genome replication
Response to virus
Reg. of viral life cycle
Cytokine-mediated signaling pathway
Innate immune response
Defense response to other organism
Cellular response to cytokine stimulus
Response to cytokine
Response to biotic stimulus
Response to other organism
KO
+TREX1
0
5
10
15
20
Fold enrichment
C
KO infected vs. mock
Neg. reg. of viral genome replication
Type I interferon signaling pathway
Cellular response to type I interferon
Defense response to virus
Reg. of viral life cycle
Response to virus
Reg. of response to biotic stimulus
Cytokine-mediated signaling pathway
Reg. of defense response
Innate immune response
Cellular response to cytokine stimulus
Response to cytokine
Defense response to other organism
Reg. of response to external stimulus
Response to other organism
Response to external biotic stimulus
Reg. of immune response
N. of Genes
10
20
30
40
-log10(FDR)
10
12
14
16
18
0
10
20
Fold enrichment
+TREX1 infected vs. mock
Defense response to protozoan
Response to protozoan
Fibrinolysis
Pos. reg. of interleukin-1 beta production
Interleukin-1 beta production
Cellular response to interferon-gamma
Defense response to virus
Response to interferon-gamma
Response to bacterium
Pos. reg. of cytokine production
Defense response to other organism
Response to biotic stimulus
Response to other organism
Response to external biotic stimulus
Innate immune response
Reg. of response to external stimulus
N. of Genes
2.5
5.0
7.5
10.0
12.5
-log10(FDR)
2.0
2.5
3.0
3.5
4.0
0
20
40
60
Fold enrichment
D
KO baseline
Neg. reg. of inflammatory response to antigenic stimulus
Reg. of inflammatory response to antigenic stimulus
Neg. reg. of inflammatory response
Inflammatory response to antigenic stimulus
Adenylate cyclase-modulating G protein-coupled receptor signaling pathway
Detection of stimulus involved in sensory perception
Detection of chemical stimulus involved in sensory perception of smell
Reg. of inflammatory response
Detection of chemical stimulus involved in sensory perception
G protein-coupled receptor signaling pathway
Sensory perception of smell
Sensory perception
Detection of chemical stimulus
Sensory perception of chemical stimulus
Detection of stimulus
Nervous system proc.
Inflammatory response
Reg. of ion transport
Trans-synaptic signaling
Ion transport
N. of Genes
50
100
150
200
-log10(FDR)
10
15
20
0
1
2
3
Fold Enrichment
+TREX1 baseline
Induction of bacterial agglutination
Nitric oxide mediated signal transduction
Homophilic cell adhesion via plasma membrane adhesion molecules
Cell-cell adhesion via plasma-membrane adhesion molecules
Sodium ion transmembrane transport
Sodium ion transport
Reg. of metal ion transport
Circulatory system proc.
Blood circulation
Cation transmembrane transport
Inorganic cation transmembrane transport
Inorganic ion transmembrane transport
Cell-cell adhesion
Cation transport
Metal ion transport
Ion transmembrane transport
Ion transport
Cell adhesion
Biological adhesion
Transmembrane transport
N. of Genes
30
60
90
-log10(FDR)
2.0
2.5
3.0
3.5
0
5
10
15
Fold Enrichment

**Figure S6 TREX1 modulates host gene expression but does not alter viral polymerase activity**

**A,** Influenza polymerase activity was measured in a mini-replicon assay in the presence of exogenous GFP-TREX1 or vector control. Data are shown as means of 3 replicates ± SEM. Pairwise T-tests tests were performed (ns = not significant).

**B-C,** Loss of TREX1 amplifies innate immune responses. **B,** Gene enrichment analysis of all host genes upregulated >4-fold during infection in TREX1-KO or complemented cells compared to their matched mock-treated cells. GO biological processes are shown. Enrichment FDR $<10^{-7}$ for all biological categories. **C,** Gene enrichment analysis of ISGs plotted in Fig 6F. This includes only ISGs induced ≥2-fold during infection in TREX1 KO cells whose induction levels change by at least 50% different between cell lines. Significantly enriched biological processes are shown along with their enrichment values.

**D,** TREX1-KO cells exhibit a chronic inflammatory state. Genes differentially expressed at least 4-fold between uninfected TREX1-KO and complemented cells were subject to gene enrichment analysis. Significantly enriched biological processes are shown along with their enrichment values.

PR8 TRPPC NS (plus-sense)

UTR
NS1
**mutant splice acceptor**
miR124
**approximate Drosh processing sites (3' of marked nt) (PMID20841420)**
sgRNA 2.0 (*PMID 25494202)*
*Shechner sgRNA target sequence (PMID 26030444)*
MS2 hairpins
NEP (5' exon in NS1 coding sequence
UTR

agcaaaagcagggtgacaaaaacataatggatccaaacactgtgtcaagctttcaggtagattgctttctttggcatgtccgcaaacga
gttgcagaccaagaactaggcgatgccccattccttgatcggcttcgccgagatcagaaatccctaagaggaaggggcagtactctcgg
tctggacatcaagacagccacacgtgctggaaagcagatagtggagcggattctgaaagaagaatccgatgaggcacttaaaatgacca
tggcctctgtacctgcgtcgcgttacctaactgacatgactcttgaggaaatgtcaagggactggtccatgctcatacccaagcagaaa
gtggcaggccctctttgtatcagaatggaccaggcgatcatggataagaacatcatactgaaagcgaacttcagtgtgatttttgaccg
gctggagactctaatattgctaagggctttcaccgaagagggagcaattgttggcgaaatttcaccattgccttctctc**ccgg**gacata
ctgctgaggatgtcaaaaatgcagttggagtcctcatcggaggacttgaatggaatgataacacagttcgagtctctgaaactctacag
agattcgcttggagaagcagtaatgagaatgggagacctccactcactccaaaacagaaacgagaaatggcgggaacaattaggtcaga
agtttgaggtaccGGGCAGGGAGAAAATTATAGTAATAGTTGCAATGAGTCACTTGCTTCTAGATCAAGATCAGAG**A**CTCTGCTCTCCG
TGTTCACAGCGGACCTTGATTTAATGTCATACAATTAAGGCACGCGGTGAATGCCAAGAGCGGA**G**CCTACGGCTGCACTTGAAGGACAT
CCGAGAGAAGTTAGGAAGGGTGGGGAGAAACAATTCTAGAATGActcgag*ATCTAGATACGACTCACTAT*GTTTTAGAGCTAGGCCAAC
ATGAGGATCACCCATGTCTGCAGGGCCTAGCAAGTTAAAATAAGGCTAGTCCGTTATCAACTTGGCCAACATGAGGATCACCCATGTCT
GCAGGGCCAAGTGGCACCGAGTCGGTGCTTTTTgcggccgcgatcaccattgccttctcttccaggacatactgctgaggatgtcaaaa
atgcagttggagtcctcatcggaggacttgaatggaatgataacacagttcgagtctctgaaactctacagagattcgcttggagaagc
agtaatgagaatgggagacctccactcactccaaaacagaaacgagaaatggcgggaacaattaggtcagaagtttgaagaaataagat
ggttgattgaagaagtgagacacaaactgaagataacagagaatagttttgagcaaataacatttatgcaagccttacatctattgctt
gaagtggagcaagagataagaactttctcgtttcagcttatttagtactaaaaacacccttgtttctact

**Figure S7 Split-NS TRPPC cassette.** Genomic sequence of the engineered NS segment.
