## Supplemental Table Legends for "Pathogen-driven CRISPR screens identify TREX1 as a regulator of DNA self-sensing during influenza virus infection"

**Supplemental Table 1: Enrichment following TRPPCa selection.**

**A-C,** UMI counts, frequency, and enrichment of individual TRPPC viruses/sgRNAs in three independent genome-wide screen.

**D,** Overlap of genes enriched >4-fold after 5 passages for Screens A-C. Fold enrichment is shown for each virus/sgRNA (i.e. VIRUS list). Enrichment scores for sgRNAs that do not meet the enrichment threshold in a specific screen are left blank, even though they might show enrichment <4-fold in the full data set. sgRNAs are collapsed to specific genes (i.e. GENE list) prior to overlap comparison.

**Supplemental Table 2: sgRNA sequences encoded by viruses in the pilot screen.**

**Supplemental Table 3: Sequence of key primers used in methods.**

**Supplemental Table 4: Master sheet of raw data and uncropped blots for Figures 1-6.**

Data for each panel is on a separate tab.

**Supplemental Table 5: Master sheet of raw data and uncropped blots for Figures S1-6.**

Data for each panel is on a separate tab.
